# Accumulation of Treg cells is detrimental in late-onset (aged) mouse model of multiple sclerosis

**DOI:** 10.1101/2021.12.16.472986

**Authors:** Weikan Wang, Rachel Thomas, Jiyoung Oh, Dong-Ming Su

## Abstract

Although typically associated with onset in young adults, multiple sclerosis (MS) also attacks aged people, which is termed late-onset MS. The disease can be recapitulated and studied in the aged mouse model of experimental autoimmune encephalomyelitis (EAE). The onset of induced EAE is delayed in aged mice, but the disease severity is increased relative to standard EAE in young mice. Given that CD4^+^FoxP3^+^ regulatory T (Treg) cells play an ameliorative role in MS/EAE severity and the aged immune system accumulates Treg cells, failure of these cells to prevent or ameliorate EAE disease is enigmatic. When analyzing the distribution of Treg cells in EAE mice, the aged mice exhibited a higher proportion of polyclonal(pan) Treg cells and a lower proportion of antigen-specific-Treg cells in their periphery, but lower proportions of pan- and antigen-specific-Treg cells in the central nervous system (CNS). Furthermore, in the aged CNS, Treg cells exhibited a higher plasticity and T effector (Teff) cells exhibited a greater clonal expansion, which disrupted the Treg/Teff balance. Transiently inhibiting FoxP3 expression in peripheral Treg cells partially ameliorated the disease and corrected Treg distribution in the aged mice. These results provide evidence that accumulated aged Treg cells play a detrimental role in neuronal inflammation of aged MS.

**Highlights:** *Question:* CD4^+^ regulatory T (Treg) cells typically play an ameliorative role in multiple sclerosis (MS) onset and severity. However, why aged immune system has accumulated peripheral Treg cells, but the elderly has more severe MS symptoms?

*Findings:* Aged Treg cells cannot easily distribute to the CNS of aged EAE mice, and those aged Treg cells that did enter the CNS exhibited increased plastic features. However, transient inhibition of peripherally accumulated Treg cells corrected Treg distribution and partially ameliorated the disease in the aged mice.

*Conclusion and mechanistic insights:* Accumulated aged Treg cells within an “inflammaging” condition do not play an ameliorative role but are potentially detrimental for inflamed CNS repair processes in aged EAE mice due to impeding the trafficking of immune cells into the inflamed CNS. 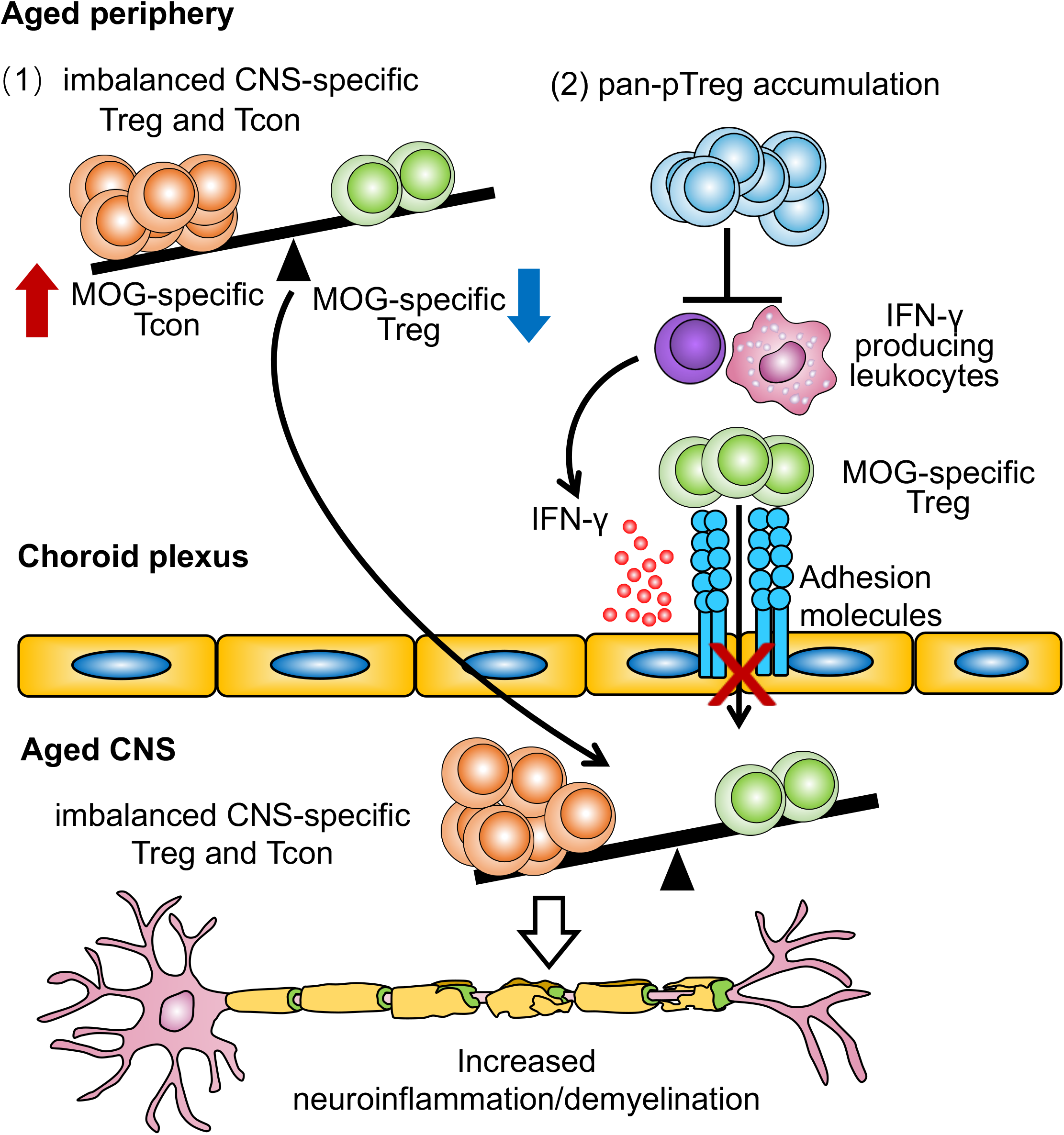

## INTRODUCTION

Neuronal inflammatory diseases are commonly seen in the elderly. These diseases are tightly associated with aberrant immune cell function ^1, 2^. Human multiple sclerosis (MS) is an autoimmune neuronal inflammatory disease, and its pathology is mainly associated with CD4^+^ T effector (Teff) cell-mediated autoimmune demyelination ^3^. Although it typically presents with onset in young adults, it either persists in a relapsing-remitting manner throughout the life of the patients, or it develops in the elderly (termed: late-onset MS) ^4, 5, 6, 7^. MS in the elderly exhibits a severe progression mediated by the aged immune milieu ^8, 9, 10^. To study MS, the mouse model of experimental autoimmune encephalomyelitis (EAE) has been widely used. However, most studies of this disease, either human MS or mouse model EAE, have focused on young subjects and have thus overlooked the age-associated T cell immune conditions. Therefore, there is insufficient knowledge regarding how MS/EAE is impacted by aged T cells, particularly aged CD4^+^ Teff and aged CD4^+^FoxP3^+^ regulatory T (Treg) cells.

T cell-mediated self-tolerance for controlling autoimmunity ^11^, including neuronal autoimmunity, is established and maintained through two primary mechanisms ^12^: depleting self-reactive T cell clones via thymocyte negative selection ^13, 14^ and suppressing aberrant immune reaction in the periphery via Treg cells ^15, 16^, which are mainly generated by the thymus ^17, 18, 19^. The thymus undergoes a progressive age-related involution during aging, which perturbs negative selection, resulting in an increased output of self-reactive Teff cells to the periphery. These self-reactive Teff cells attack self-tissues inducing inflammation, thereby enhancing senescent somatic cell secretions which underlie chronic inflammation (termed senescence-associated secretory phenotype) ^20, 21, 22^ in the elderly (termed inflammaging) ^23^. Meanwhile, the aged thymus exhibits relatively enhanced thymic Treg (tTreg) generation ^24^ which join the accumulated peripheral Treg (pTreg) pool in the aged periphery ^25, 26^. This accumulation of polyclonal (pan)-pTreg cells is disadvantageous for anti-infection ^27^ and anti-tumor ^28, 29, 30^ immunity, as well as vaccination efficacy in the elderly ^31^. In elderly patients with neurodegenerative Alzheimer’s disease (AD) or Parkinson’s disease, the frequency of peripheral Treg cells and their expression of FoxP3 are increased. However, they do not help to ameliorate these diseases, but rather are correlated with increased disease severity ^32^.

Given that Treg cells generally play an ameliorative role in MS disease onset and severity ^33, 34, 35^ and the aged immune system exhibits accumulated pan-pTreg cells, we asked why these cells do not ameliorate severity of MS in elderly patients. Mounting evidence shows that patients over the age of 65 are more likely to have a severe progressive course of MS. Primary progressive (PPMS, disease without remissions) and secondary progressive (SPMS, irreversible damage and disability) are severe and commonly seen in aged patients. However, aged patients are less likely to have a mild course, such as relapsing-remitting MS (RRMS) (<40%), compared to their young counterparts of whom >80% having RRMS ^5, 36^. We proposed that the answer to this question is a dichotomy of the Treg cells’ role in protection from CNS inflammation within the aged immune system ^1, 2^. We believe that Treg cells have either a detrimental or beneficial role, which is dependent on their location, either inside the CNS or outside the CNS (in the periphery). In addition, the imbalance between Treg and Teff cells ^37^ and Treg functional plasticity ^38^ within the aged CNS could determine disease severity.

To verify our hypothesis, we investigated how accumulated aged Treg cells impact late-onset MS using the EAE model in aged mice, by examining Treg distribution inside and outside the CNS, Treg cell antigen specificity to myelin oligodendrocyte glycoprotein (MOG) and pan-antigens, and Treg function-associated molecular profiles during EAE disease in young and aged animals. We also transiently inhibited FoxP3 expression in accumulated pan-pTreg cells in the aged mice and demonstrated a partial amelioration of the disease in the aged mice. Finally, we verified the proposed underlying mechanism and obtained mechanistic insights that accumulated Treg cells residing at a CNS barrier membrane, the choroid plexus (CP), potentially impede the trafficking of immune cells into the inflamed CNS. This results in disruption of the balance between Treg and Teff cells, thereby inhibiting Treg suppression of Teff cell clonal expansion in the inflamed CNS. In addition, CNS-infiltrated Treg cells also showed increased plasticity exhibited by co-expression of IFN-g and/or IL-17A along with FoxP3, which results in reduction of suppressive capacity. Together, our results provide novel evidence that accumulated and compromised aged Treg cells do not play an ameliorative role but are potentially detrimental during late-onset (aged) neuronal autoimmune EAE disease course and severity.

## RESULTS

### The course of late-onset EAE disease of aged mice exhibited distinct characteristics

Since T cell-mediated autoimmune MS disease typically has an onset in young adults, studies on MS etiology, pathology, immunology, disease courses, etc., have been focused on young patients (20~40 years old) or have used an EAE model with young mice (2-3 months old). However, late-onset (aged) MS in the elderly (diagnosed at the age of 60 or over) has been reported ^4, 5, 6, 7^. More information about late-onset MS remains to be determined. Therefore, we should develop a late-onset EAE mouse (≥18 months of age) mouse model.

Firstly, we used a standard immunization protocol (Hooke Lab etc.) ^39^, i.e. immunized the same dosage of MOG peptide and pertussis toxin (PT) to each young and aged C57BL/6 wild-type (WT) female mice (i.e. per mouse based dosage, Fig. S1), which was widely adopted in previous studies. We found that the aged female mice have dichotomous of EAE courses. Some aged mice (about 1/3) had delayed EAE onset but their symptoms became severe after onset, compared to the young group (Fig. S1B, Type-I course), whereas some other aged mice (about 2/3) showed lower debility than the young group and never reached debility score > 3.0 by 48 days after the first immunization with MOG. Then, we gave them, along with similar numbers of young controls which received the first immunization, a second immunization (Fig. S1A). Interestingly, these aged mice with lower debility immediately showed increased EAE symptoms and became much more severe than their young counterparts. We termed this Type-II course (Fig. S1C).

As the body weight is a critical factor for the dosage administration of drugs, and the aged mice (30~40g) have about 50% more to double body weight than the young (20~25g) ones (information from our colonies and the Jackson laboratory: https://www.jax.org/jax-mice-and-services/strain-data-sheet-pages/body-weight-chart-000664#, and https://www.jax.org/jax-mice-and-services/strain-data-sheet-pages/body-weight-chart-aged-b6.). We believe that the inconsistent EAE progression/severity in the age mice might not be due to the inconsistent susceptibilities compared to the young mice, but due to the insufficient MOG_35-55_ peptide administration. Therefore, we adjusted the MOG_35-55_ peptide at a body weight dependent manner (80μg/10g body weight) and the dosage of PT (100ng/10g body weight) as in Fig. 1A. Then, we found that young and aged mice have a consistent EAE progression, despite different severities (Figs. 1B and C). Therefore, we, for the first time, established a late-onset (aged) EAE model on C57BL/6 WT mice with a body weight corrected MOG_35-55_ peptide dosage immunization.

**Figure 1.**
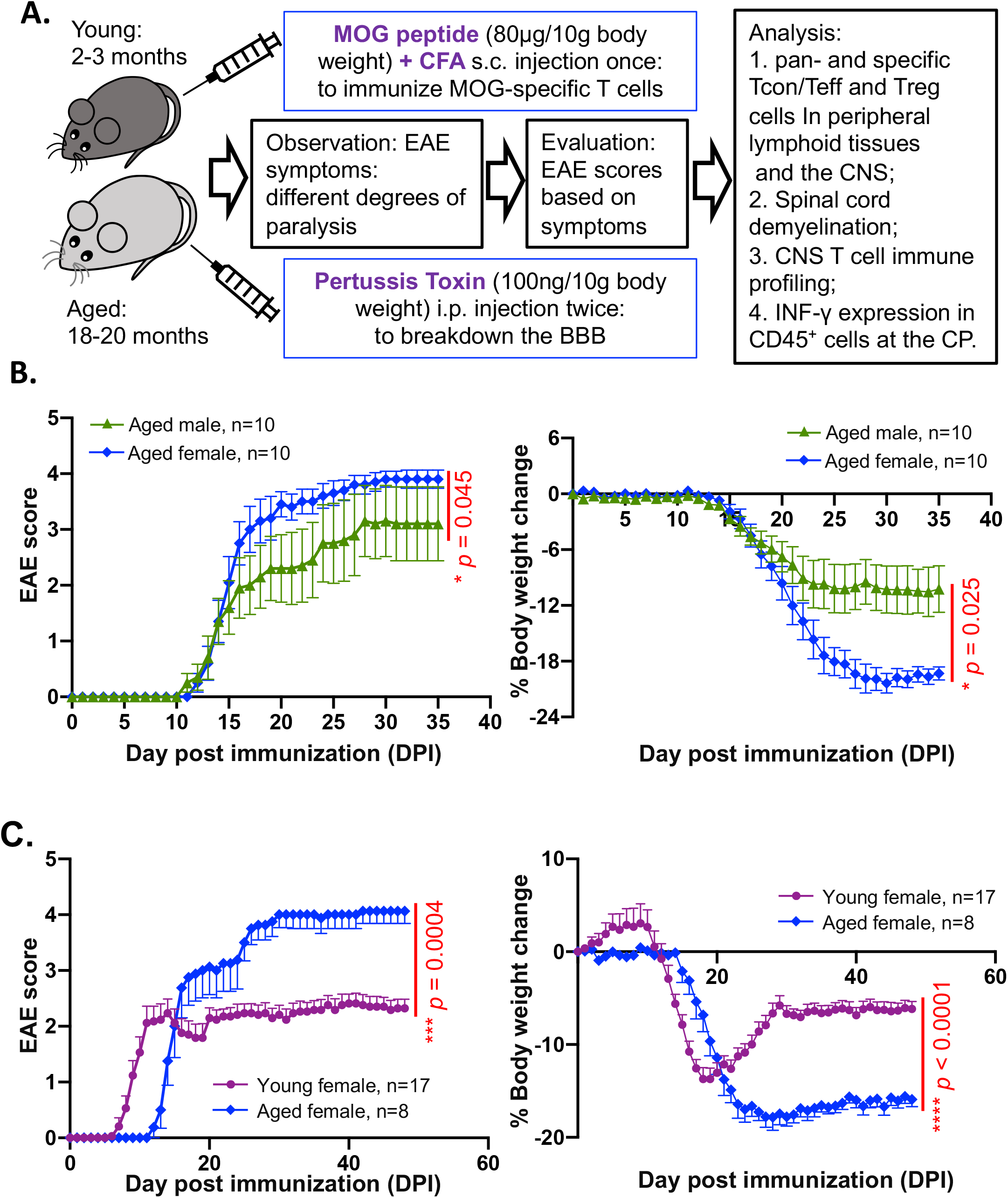
Characteristics of EAE disease in young versus aged mice. **(A)** Workflow for immunization and induction of EAE in C57BL/6 mice with reduced dose (80μg/10g body weight) of MOG peptide and follow-up analysis. **(B)** The variability of EAE pathological scores (left panel) and body weights during disease onset (right panel) in aged male and female mice. **(C)** Characteristics of EAE pathological scores (left panel) and body weights during disease onset (right panel) in young (cherry-color line) versus aged (blue line) female mice. The *p*-values of EAE scores were calculated by Mann-Whitney *U* test and body weight changes were calculated by two-way repeated-measures ANOVA with Geisser-Greenhouse correction, and a statistically significant difference was considered to be *p < 0.05.*

Next, we investigated disease progression with the adjusted dosage protocol (Workflow shown in Fig. 1A). We first compared the courses of EAE disease in aged male and female mice (Fig. 1B) and found the two groups had similar EAE onset on 11 or 12-days post-immunization (DPI). However, the male group had less severity and great variation of disease symptoms (Fig. 1B, green lines). In contrast, female mice had minimal variation in the disease course with severer disease symptoms (Fig. 1B, blue lines). These characteristics are consistent with observations in human MS disease, in which women are the predominant population of MS patients, and are also consistent with most published reports using female mice for EAE research ^41, 42^. Therefore, we used female mice for the rest of our study. By comparing young and aged female mice, we determined that aged mice usually had a distinct EAE onset course, i.e. late-onset course. Aged mice had an EAE onset about 12-DPI, which was about 6-day later than EAE onset in young mice (as early as at 6-DPI); however, aged mice developed a more severe disease course (Fig. 1C, blue lines, demonstrated by either EAE scores: left panel, or loss of body weight: right panel). The results provide novel evidence that pathological severity and course of EAE disease in young and old individuals are distinctively different. Although aged mice had a delayed EAE onset after immunization, they suffered from more severe disease progression compared to young mice.

### Different distributions of Treg cells between young and aged mice during EAE disease

Given that Treg cells play a vital protective role in the regulation of MS/EAE severity and progressive disease course ^35, 40, 41, 42, 43, 44, 45^, we compared the distributions of pan- and MOG-specific Treg cells, along with Teff (or termed conventional T, Tcon) cells, in the periphery (secondary lymphoid organs, Fig. 2) and the CNS (brain and spinal cord, Fig. 3) between young and aged mice during EAE disease (gating strategies were shown in Fig. S2). We indeed found similarities and differences in the distributions between young and old, which hint at the possibility that Treg cells are involved in the characteristics of EAE disease course.

**Figure 2.**
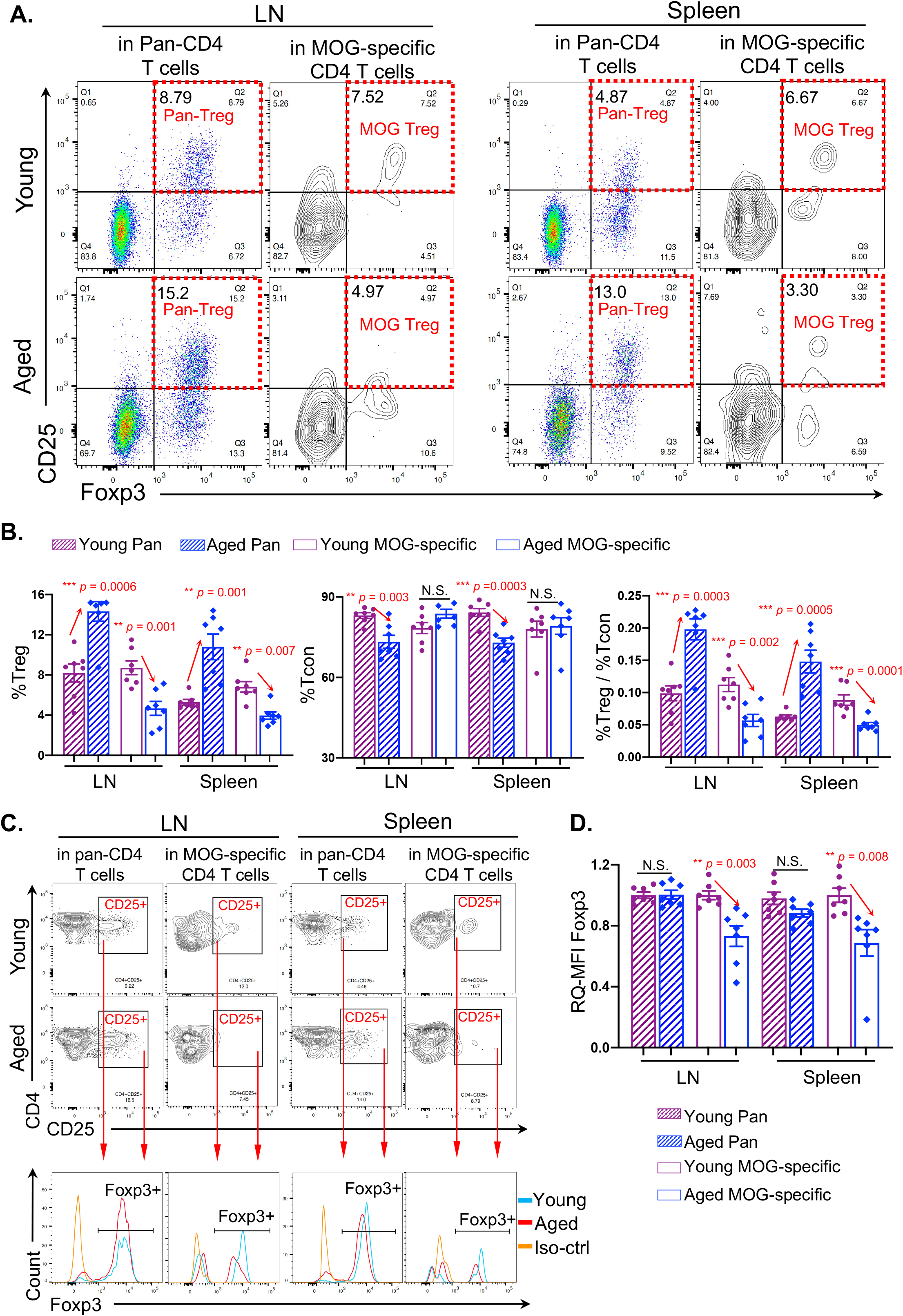
Imbalanced distributions of pan- and MOG-specific Treg cells in the periphery of late-onset EAE disease of aged mice. Mice were immunized as Fig. 1A workflow. Then, T cells in LNs and spleen were analyzed. **(A)** Flow cytometry gating strategies show pan (poly-clonal)- and MOG-specific Treg cells in LNs and spleens of young and aged mice. MOG-specific gate was determined by a dot-plot of MHC-II MOGspecific I-A^b^ tetramer versus MHC-II control tetramer I-A^b^. **(B)** Summarized results of the percentages of pan- (striped bars) and MOG-specific (open bars) Treg cells (left panel) and Teff cells (middle panel), and ratios of Treg/Teff cells (right panel) in LNs and spleen between young (cherry) and aged (blue) mice. **(C)** Flow cytometry gating strategies show FoxP3^+^ peaks (bottom panel) from CD4^+^CD25^+^ gates of young (top panel) and aged (middle panel) mice. **(D)** Summarized results of the relative quantitative (RQ) mean fluorescent intensity (MFI) of FoxP3 expression in pan- (striped bars) and MOG-specific (open bars) CD4^+^CD25^+^ population in LNs and spleen between young (cherry) and aged (blue) mice. In panels B and D, each symbol represents an individual animal sample. The *p*-values between two groups were analyzed by unpaired *Student’s t-test,* and a statistically significant difference was considered to be *p < 0.05.* “N.S.” stands for “not significant”, and error bars indicate mean ± SEM.

**Figure 3.**
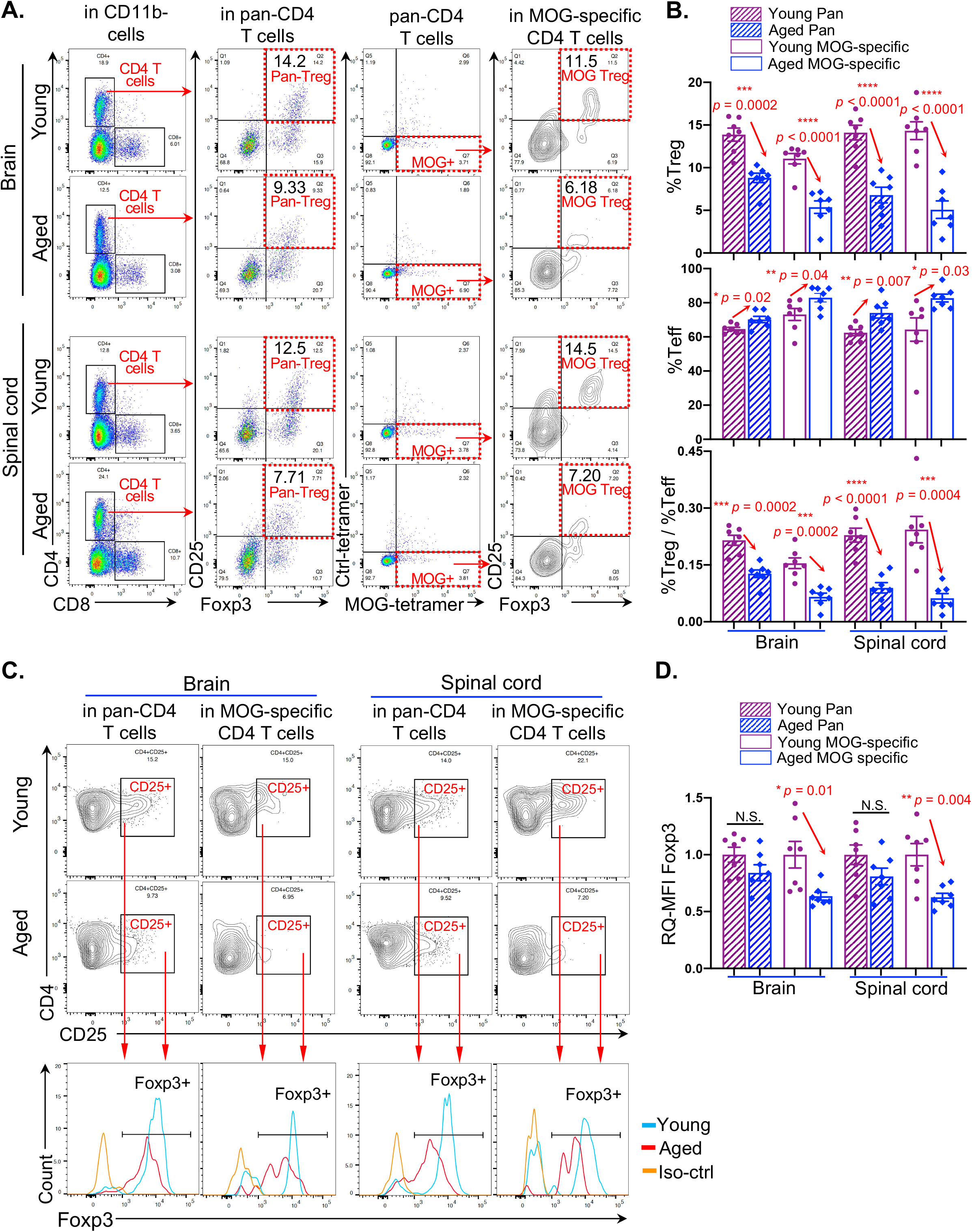
Imbalanced distributions of pan- and MOG-specific Treg in the CNS of late-onset EAE disease of aged mice. Mice were immunized per Fig. 1A workflow. Then, T cells in the CNS were isolated via gradient centrifuge and analyzed by flow cytometry. **(A)** Gating strategies show pan-Treg population (left two columns) and MOG-specific Treg population (right two columns) in the CNS. **(B)** Summarized results of the percentages of pan- (striped bars) and MOG-specific (open bars) Treg cells (top panel) and Teff cells (middle panel), and ratios of Treg/Teff cells (bottom panel) in the CNS (brain and spinal cord) between young (cherry) and aged (blue) mice. **(C)** Flow cytometry gating strategies show FoxP3^+^ peaks (bottom panels) from CD4^+^CD25^+^ gates of the CNS (brain and spinal cord) of young (top panel) and aged (middle panel) mice. **(D)** Summarized results of the relative quantitative (RQ) mean fluorescence intensity (MFI) of FoxP3 expression in pan- (striped bars) and MOG-specific (open bars) CD4^+^CD25^+^ populations in the brain and spinal cord between young (cherry) and aged (blue) mice. In panels B and D, each symbol represents an individual animal sample. The *p*-values between two groups were analyzed by unpaired *Student’s t-test,* and a statistically significant difference was considered to be *p < 0.05,* “N.S.” stands for “not significant”, and error bars indicate mean ± SEM.

In the periphery, either in the lymph node (LN) or spleen, the percentage of pan-Treg cells in aged EAE mice was increased (Figs. 2A and B stripped bars in left and right panels). This is consistent with published reports regarding the accumulation of pTreg cells in aged individuals ^25, 46, 47, 48^. However, the percentage of MOG-specific Treg cells in aged EAE mice was decreased (Figs. 2A and B, opened bars in left and right panels). The expression of FoxP3, which is tightly associated with Treg suppressive function, was decreased in MOG-specific Treg cells of aged EAE mice (Figs. 2C and D). In addition, pan-Teff cells were decreased (Fig. 2B, striped bars in middle panel), but MOG-specific Teff cells were not changed in aged EAE mice (Figs. 2B, open bars in middle panel). The results suggest that although pan-Treg cells are increased, MOG-specific Treg cells are reduced. This, coupled with unchanged MOG-specific Teff cells in the aged periphery, results in an imbalance of MOG-specific immunopathogenic and immunoregulatory T cells during aged EAE onset.

In the CNS, including the brain and spinal cord, the percentages of both pan-Treg and MOG-specific Treg cells were decreased in aged EAE mice (Figs. 3A and B top and bottom panels). The results suggest that peripheral Treg cells could not enter the aged, inflamed CNS. The expression of FoxP3 in MOG-specific Treg cells within the aged, inflamed CNS was decreased (Figs. 3C and D), similar to those in the periphery (Figs. 2C and D), suggesting that even if the MOG-specific Treg cells enter the aged CNS, they cannot fully execute their suppressive function. Particularly, both pan- and MOG-specific Teff cells were increased in aged EAE mice (Fig. 3B middle panel), different from those in the periphery (Fig. 2B middle panel), suggesting that both CNS-infiltrated pan- and MOG-specific Teff cells interact with the CNS tissues to participate in inflammation. However, both pan- and MOG-specific Treg cells are less robust in controlling the inflammation in the CNS of aged EAE disease. Interestingly, such a discrepancy between the young and aged mice Treg cells was not observed prior to EAE onset (Fig. S3), indicating that the imbalanced Treg distribution in the aged mice developed during EAE disease.

Together, these findings can help explain one of the reasons that aged mice have a late-onset but more severe EAE pathology. Delayed onset is probably due to the pTreg accumulation, while the more severe progressive disease course is perhaps due to impaired trafficking of both pan- and MOG-specific Treg cells into the CNS during EAE disease. In other words, although pan-pTreg cells are accumulated in the periphery of aged mice, it is likely that they cannot easily enter into the aged, inflamed CNS during EAE disease.

### CNS-infiltrated Treg cells in late-onset EAE of aged mice exhibited dysfunctional molecular profiles

Treg cells can possess relatively unstable features ^49, 50^, including down-regulation of FoxP3 expression ^16, 51^ as confirmed in our aged late-onset EAE mouse model (Figs. 2D, 3D), and co-expression of *ifng* or *il17a* toward Th1-like or Th17-like plastic conversion ^52, 53^ upon autoimmune stimulation ^16, 51, 54^ This leads to functional plasticity, resulting in increased pathology and reduced suppressive capacity ^16, 53^. We believe that this unstable phenotype will likely be more prominent in the aged microenvironment. To evaluate these function-related molecular profiles, we analyzed the expression of *ifng* and *il17a* genes in CNS-infiltrated Treg cells of EAE mice at the single-cell level (Fig. 4A). We compared CD4^+^FoxP3^+^ Treg cells in the young and aged CNS (Fig. 4B) during EAE disease and found that expression of *ifng* and *il17a* was indeed increased in the CNS-infiltrated aged Treg cells (Fig. 4C). In addition, we noticed that CD8^+^ T cells in the CNS ^55^, particularly, activated or pathogenic CD8^+^ T cells, indicated by expressions of *ifng* and *il17a,* which potentially lead to autoimmune encephalomyelitis ^57, 58^, presented increased *ifng* and *il17a* expressions in the aged CNS with EAE disease (Fig. 4D). IL-17A secreting CD8^+^ T cells, termed Tc17 cells, in the CNS support Th17-mediated autoimmune encephalomyelitis ^56, 57^. These results imply that aged Treg cells, which have infiltrated into the CNS after EAE onset, could have reduced capacity to suppress neuronal inflammation.

**Figure 4.**
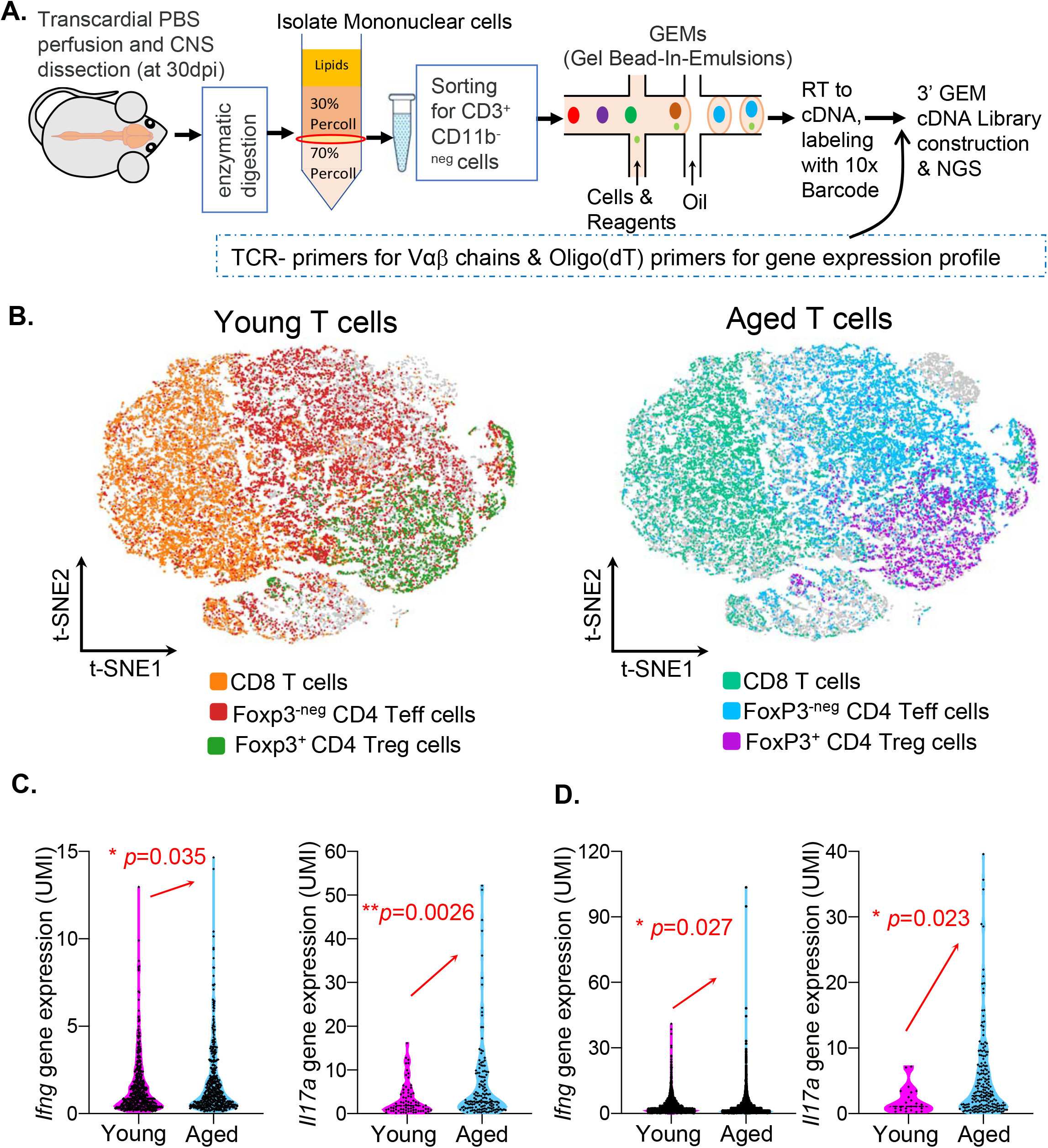
Treg function-associated profile analysis of altered Treg plasticity and pro-inflammatory CD8 T cells in the CNS of aged mice during late-onset EAE. Mice were immunized per Fig. 1A workflow. **(A)** Workflow of sc-RNA-seq for CNS infiltrated T cells. T cells in the CNS (brain and spinal cord) were isolated via gradient centrifugation and sorted for CD3^+^CD11b^-neg^ cells by flow cytometry 30 days after immunization. Then, single cells were captured and emulsified with specific gel beads for reverse transcription and construction of cDNA library for high-throughput sequencing. **(B)** The pattern of young (left panel) and aged (right panel) T cells from the CNS of three young and three aged EAE mice with t-distributed stochastic neighbor embedding (t-SNE) plots. **(C)** Expressions (Unique Molecular Identifier, UMI) of *ifng* (left panel) and *il17a* (right panel) genes in single Treg (CD4^+^FoxP3^+^) cells of the young (cherry) and aged (blue) CNS. **(D)** Expressions of *ifng* gene (left panel) and *il17a* (right panel) in single CD8^+^ T cells of the young (cherry) and aged (blue) CNS. Data in panels C and D were analyzed by unpaired *Student’s t-test,* and a statistically significant difference was considered to be *p < 0.05.* Each symbol represents a single cell.

### Clonal expansions of CNS-infiltrated Teff cells in late-onset EAE in aged mice was increased

To assess whether CNS-infiltrated aged Treg cells have reduced suppressive capacity, a comparison of Teff cell clonal expansions in the EAE CNS can shed some light. We therefore analyzed clonal expansions in the CNS-infiltrated Teff cell pool along with the Treg cell pool based on TCR sequence similarity via TCRαβ immune profile analysis with the single-cell RNA sequencing (sc-RNA-Seq) approach (Fig. 5). The capacity of Treg cells to suppress clonal expansions of various Teff cells influences Teff cell-induced CNS inflammation. The results showed that aged Teff cells had a greater clonal expansion, compared to the young (Fig. 5A top pies and Fig. S4 left two columns), and the expanded Teff clones (both total expanded clones and top 10 expanded clones) occupied a higher proportion of the Teff pool (Fig. 5B left panels) in the aged, inflamed CNS. However, although Treg cells can proliferate *in vivo* ^58, 59^, aged Treg clones showed the same expansion as in the young (Fig. 5A bottom pies, 5B right panels and Fig. S4 right two columns). Together, aged Teff cells are more activated in the aged, inflamed CNS during EAE disease, which is potentially due to the insufficient suppression by Treg cells. The results also indicate that the expansive capacity of CNS-infiltrated Treg cells in aged EAE mice is not reduced compared to their young counterparts.

**Figure 5.**
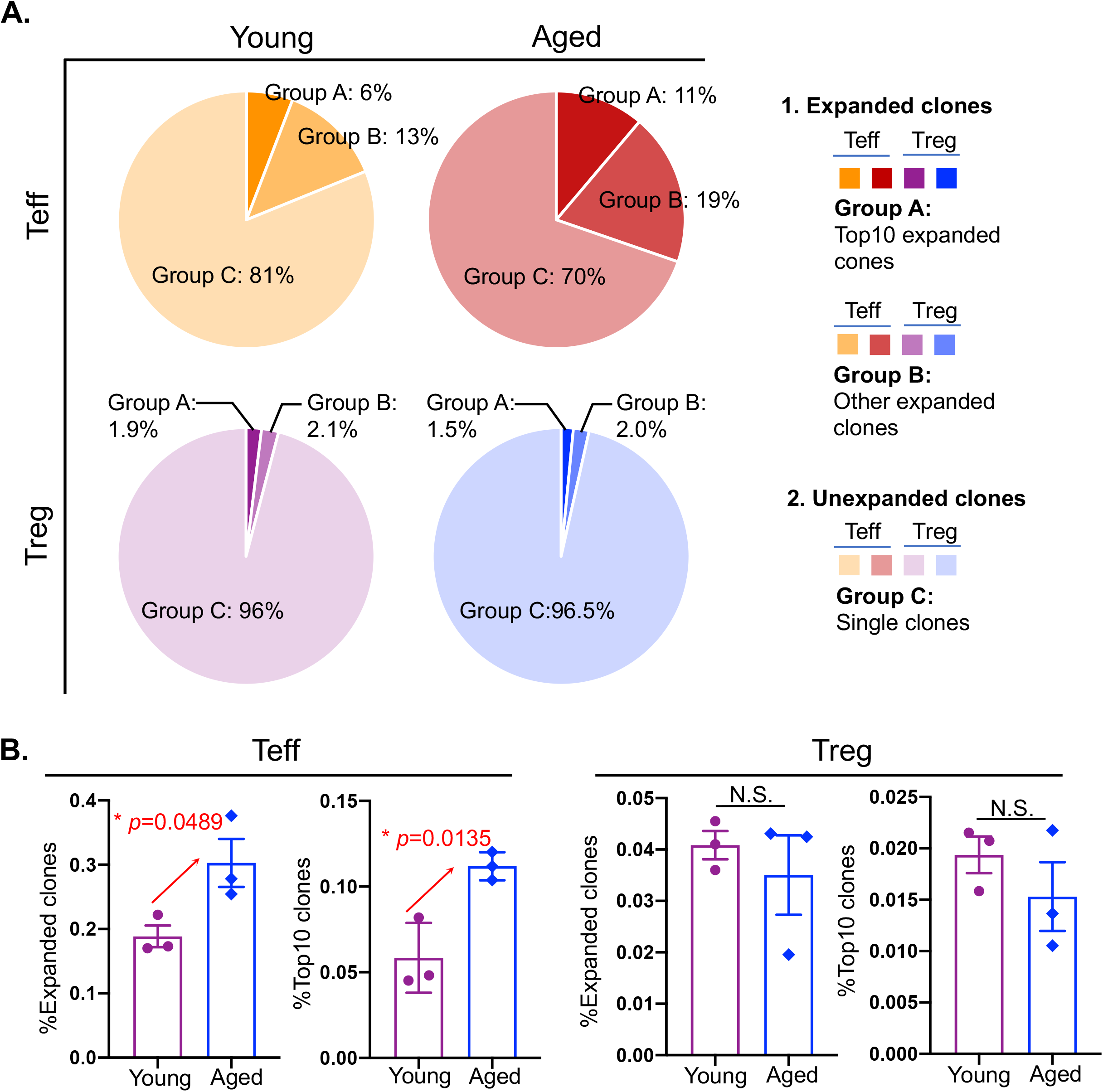
Clonal expansion in CNS-infiltrated CD4^+^ T populations of young and aged mice during EAE. Based on TCRαβ sequence similarity from single-cell immune profile RNA-Seq analysis, CNS-infiltrated Teff and Treg cells were divided into expanded T clones (more than one similar TCR sequences) and unexpanded T clones (unique TCR sequence, with clone size = 1). In the expanded T clones, we further divided them into Group-A: top 10 clones (with most similar TCR sequences) and Group-B: other expanded clones (all other ≥ 2 similar TCR sequences). **(A)** Pie charts show expanded clones (Groups-A and -B) and unexpanded clones (Group-C) of Teff (top two pies) and Treg (bottom two pies) cells in total CD4^+^ T cells from young (left two pies) and aged (right two pies) CNS during EAE. **(B)** Summarized results of the frequencies of all expanded clones of Teff (left panel) and Treg (right panel) cells in total CD4^+^ T cells from young (cherry open bars) and aged (blue open bars) CNS during EAE. Each symbol represents one mouse. Data were analyzed by unpaired *Student’s t-test*, and a statistically significant difference was considered to be *p < 0.05*.

### EAE severity was mitigated after transient inhibition of FoxP3 expression in pan-pTreg cells in the aged mice

Accumulation of pan-pTreg cells in aged individuals ^25, 46, 47, 48^ was demonstrated to be harmful during neurodegenerative disease AD, since transiently inhibiting FoxP3 expression in the accumulated Treg cells attenuated the pathology ^60^. However, it is unknown whether this accumulation is detrimental or beneficial in late-onset MS/EAE. Given that aged EAE disease occurs with accumulated Treg cells in the periphery (Fig. 2B), but less in the aged, inflamed CNS (Fig. 3B), we hypothesized that the cellular trafficking may be impeded by the accumulation outside the CNS. Therefore, transient inhibition of FoxP3 expression in accumulated pTreg cells in the aged mice could mitigate late-onset EAE symptoms and pathology. Therefore, we administrated 5-doses of P300i, which can impair Treg suppressive activities by inhibiting FoxP3 expression without affecting Teff cell responses ^61^ and has been used to transiently and partially suppress pan-pTreg cells residing outside the CNS ^60^, to aged mice beginning on 12-DPI (Fig. 6A). We evaluated the percentages of Treg cells (Fig. 6B left panel and Fig. S5A) and expression level (via mean florescence intensity, MFI) of FoxP3 (Fig. 6B right panel and Fig. S5B) in the peripheral blood at three time points (Fig. 6A three red arrowheads indicate tests, i.e. before and after the P300i injections). The results show that the pan-pTreg cells decreased one day after the last P300i injection, but returned to normal levels after 12 days (Fig. 6B). This confirms that the inhibition was effective and transient, and Treg cells had restored FoxP3 expression after the pharmaceutical inhibitory effects decay over time.

**Figure 6.**
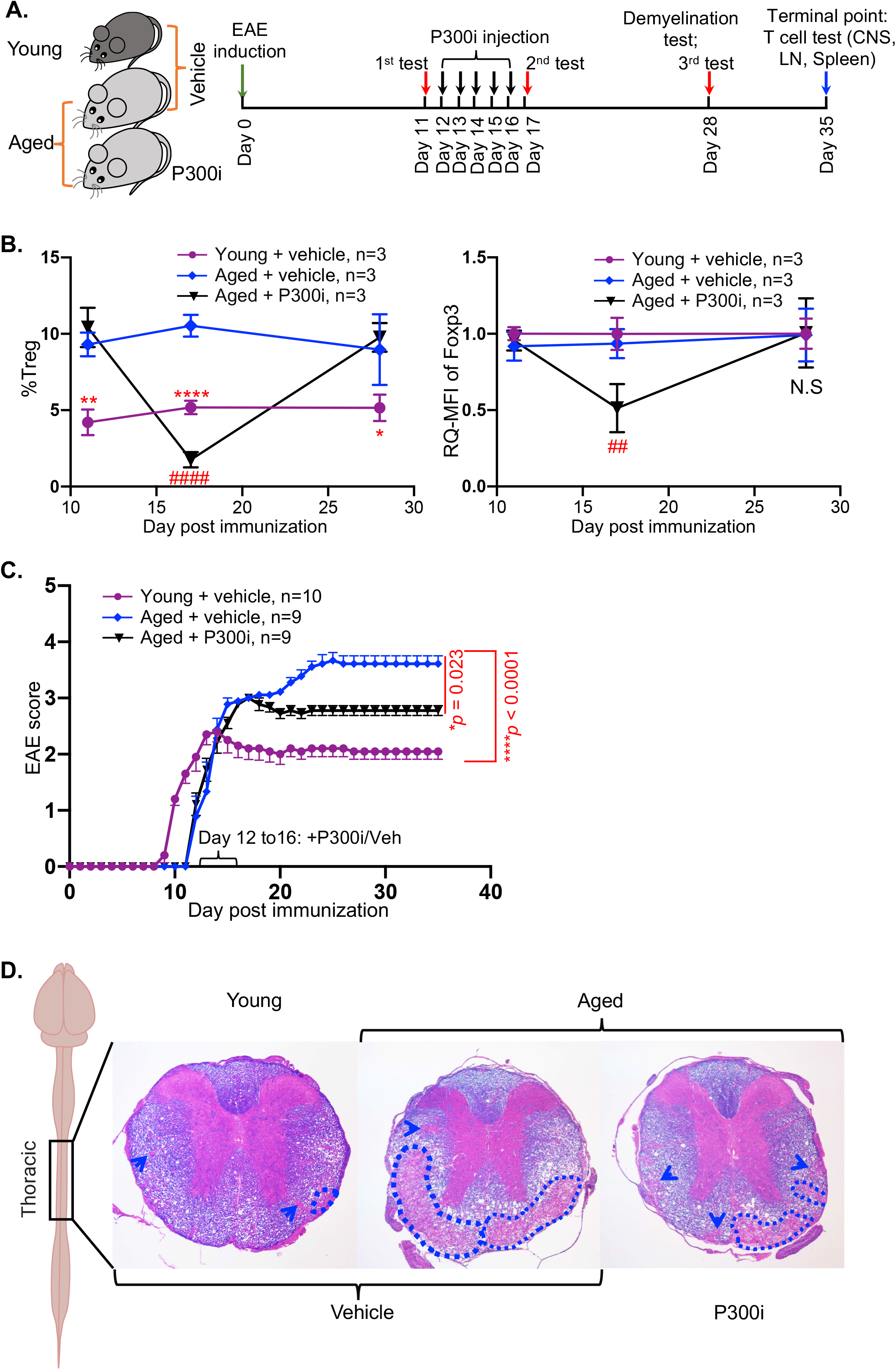
Alleviation of EAE severity after transient inhibition of FoxP3 expression in the accumulated pTreg cells in late-onset EAE in aged mice. **(A)** Workflow of EAE induction, transient inhibition of pTreg cells, blood collection time-points (red arrowheads) for Treg cell tests, and demyelination assay. **(B)** Effects of transient inhibition of pTreg cells in aged mice. At the first evaluation (one day before the first injection of P300i or vehicle), the percentages of pTreg cells in the two aged groups were higher than the young (left panel). At the second evaluation (one day after the last injection of P300i or vehicle), the percentages of pTreg cells and expression (MFI) of FoxP3 in the aged group treated with P300i (black triangles/line) were significantly reduced. At the third evaluation (12 days after the last treatment), the pTreg cells in the P300i-treated aged group were restored to the same levels as the vehicle-treated control aged group (blue dots/line). All data are expressed as mean ± SEM and were analyzed by One-way ANOVA followed by Dunnett’s multiple post-hoc test. \**p* < 0.05 young mice +vehicle v.s. aged mice +vehicle; ** *p* < 0.01 young mice +vehicle v.s. aged mice +vehicle; **** *p* < 0.0001, young mice +vehicle v.s. aged mice +vehicle; ##*p* < 0.01, aged mice + vehicle v.s. aged mice + P300i; ####*p* < 0.0001, aged mice + vehicle v.s. aged mice + P300i. Animal numbers in each group are listed in each panel. **(C)** Alleviation of the symptoms in late-onset EAE of aged mice. The EAE scores of 5x P300i-treated aged group (black triangles/line) were significantly reduced compared to their counterparts, the vehicle-treated control aged group (blue dots/line). The *p*-values were calculated using Kruskal-Wallis test followed by Dunnett’s multiple comparisons post hoc test for pairwise comparisons of groups. **(D)** Illustration of a mouse brain and spinal cord indicating the thoracic segment of the spinal cord for LFB-eosin staining (rightmost illustration). Three representative spinal cord section images of LFB-eosin staining, showing alleviation of demyelination in the spinal cords of aged mice treated with 5x P300i (right image), compared to their counterparts, the vehicle-treated age-matched control mice (middle image). The dotted outlines indicate areas of large foci of demyelination, and blue arrow heads show small foci of demyelination. This experiment was repeated three times with three mice in each group with essentially identical results.

Regarding the severity of EAE disease after transient inhibition of FoxP3 in accumulated pan-pTreg cells in the aged mice, we evaluated EAE symptoms (disease scores and demyelination) and found that the severity was attenuated in the aged P300i-treated group (Fig. 6C black line and Fig. 6D rightmost image). Although the improvement did not restore to the same level as the young group, it was significant. The results provide evidence that the accumulation of aged pan-pTreg cells in the periphery is not beneficial but rather detrimental to late-onset MS/EAE in aged individuals.

To reconfirm aforementioned results that impairing the accumulated pan-pTreg in the aged mice can rescue the EAE symptom, we introduced aged Foxp3-DTR/EGFP mice (termed DTR mice) for late-onset EAE model. DTR mice have FoxP3^+^ Treg cells expressing diphtheria toxin receptor (DTR) and thus Treg cells can be transiently depleted after administrating diphtheria toxin (DT)^62^. Therefore, we i.p. injected DT once to the aged-DTR EAE mice with at 50μg/kg body weight on 12-DPI. We evaluated the percentages of Treg cells in the peripheral blood one day before the injection (11-DPI), one day after the injection (13-DPI) and 16 days after the injection (28-DPI). As expected, peripheral blood Treg cells were completely depleted one day after the DT injection and restore to normal level 16 days after the injection (Fig. S6A and B). As a result, EAE symptom presented a partially alleviated trend in the aged-DTR mice with transient pTreg depletion (Fig. S6C).

### Transient inhibition of FoxP3 expression in the accumulated pan-pTreg cells was a potential mechanism of the late-onset EAE alleviation in aged mice

We wanted to elucidate the underlying mechanism by which late-onset EAE alleviation occurred in aged mice via transient inhibition of FoxP3 expression. Based on a published report, accumulated Treg cells, which are adherent to the barrier sites including the blood-brain barrier (BBB) and the CP of the CNS, suppress IFNg-secreting cells and potentially result in hampered trafficking of monocytes and antigen (Ag)-specific Treg cells into the inflamed CNS ^63^. Transient, rather than permanent, inhibition of FoxP3 expression in the accumulated pTreg cells mitigated AD ^60^ due to facilitated trafficking of anti-inflammatory monocytes and Treg cells during CNS inflammation. We believe this is likely the case with late-onset MS/EAE. Thus, we investigated the expression of IFN-g in CP-adherent CD45^+^ hematopoietic cells and the proportions of CNS-infiltrated Treg and Teff cells with/without the transient inhibition in the aged mice.

Once we transiently inhibited FoxP3 expression in the accumulated pan-pTreg cells in the aged mice, one day after the last treatment (Fig. 6A, the 2^nd^ test red arrow), we found that expression of IFN-g was increased in the CP-adherent hematopoietic CD45^+^ cells (Fig. 7A black line in the pick box). We also determined the impact on Treg CNS-infiltration after the pan-pTreg transient inhibition of pan-pTreg cells. The results showed that both the percentages of pan-Treg cells (Figs. 7B left panel and C black striped bar in left panel) and MOG-specific Treg cells (Figs. 7B right panel and C black open bar in left panel) were increased in the inflamed CNS of aged mice. The lack of change in FoxP3 expression in CNS-infiltrated Treg cells (Figs. 7D and E) with/without transient inhibition may indicate that the drug P300i does not affect CNS-infiltrated Treg cell function. In addition, increased Treg cells in the CNS could either suppress Teff cells (Fig. 7C middle panel) or increase the ratio of Treg versus Teff cells (Fig. 7C right panel). This ratio was previously imbalanced during late-onset EAE disease (Fig. 3B bottom panel). Likewise, this partially restored Treg to Teff balance in the aged EAE CNS was also observed in the aged-DTR EAE mice with transient pTreg depletion (Fig. S6D).

**Figure 7.**
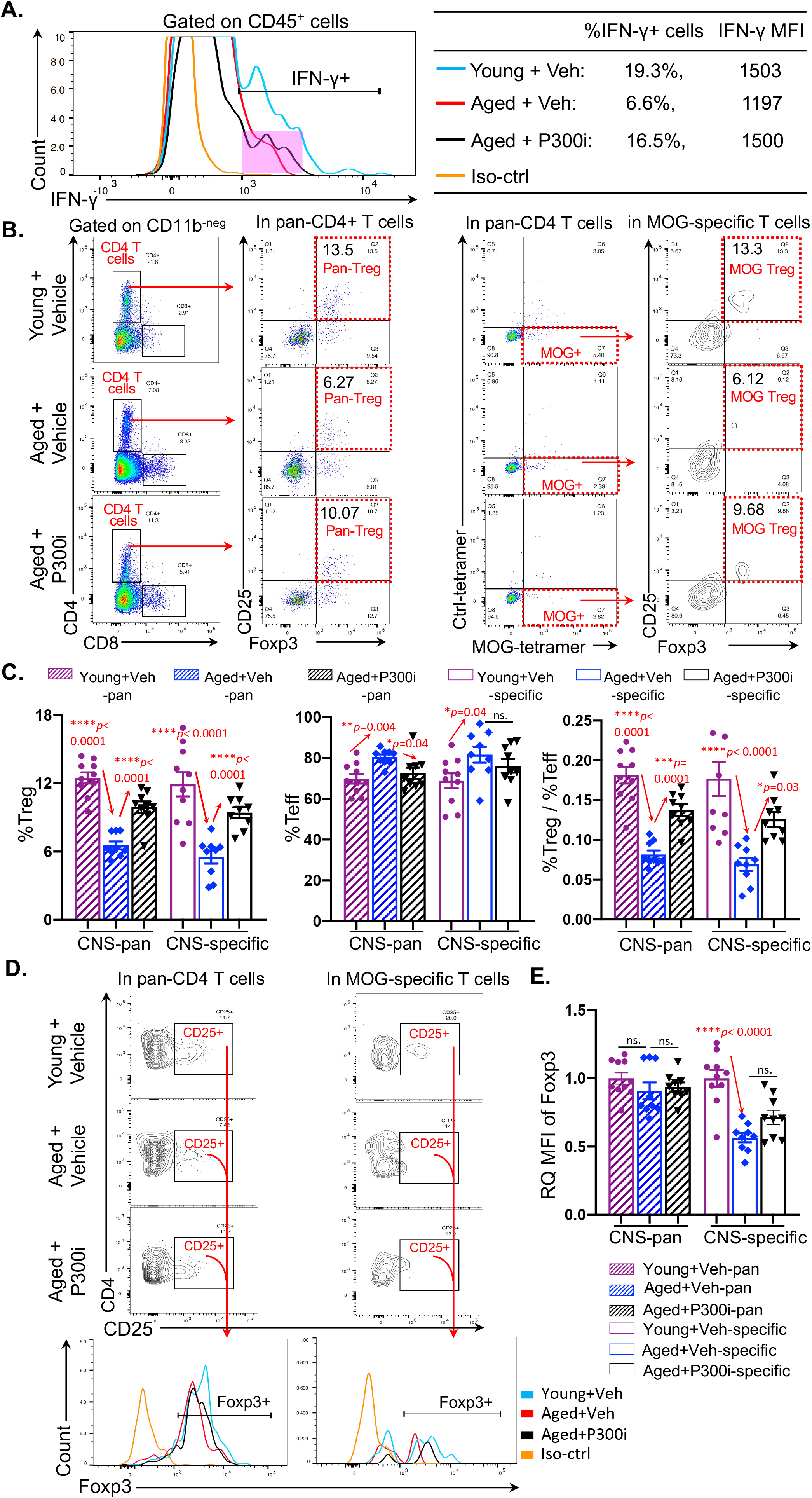
Transient inhibition of accumulated pan-pTreg cells induced the alleviation of late-onset EAE in aged mice, potentially due to restored immune cell trafficking. The workflow of the pTreg inhibition is shown in Fig. 6A. **(A)** Expression of IFN-g^+^ in CP-residing CD45^+^ cells were analyzed one day after the last P300i treatment (Fig. 6A, the second test red arrow). A representative histogram shows IFN-g^+^ CD45^+^ cells residing at the CP isolated from the brains of young and aged mice treated with P300i or vehicle. Percentages of IFN-g^+^ cells in CD45^+^ cells and MFI of IFN-g expression are listed in the table (right). **(B)** Flow cytometry gating strategies of pan-Treg and MOG-specific Treg cells, similar to Fig. 3A, in the CNSs of young and aged mice with P300i- or vehicle-treatment. **(C)** Summarized results of the percentages of pan- (striped bars) and MOG-specific (opened bars) Treg cells (left panel) and Teff cells (middle panel), and ratios of Treg/Teff cells (right panel) in the CNS (a combination of the brain and the spinal cord) among young (cherry) and aged mice, treated with vehicle (blue) or with P300i (black). **(D)** Flow cytometry gating strategies show representative FoxP3^+^ gates (bottom panels) from CD4^+^CD25^+^ gates of the CNS (brain and spinal cord) of the three groups of mice. **(E)** Summarized results of the RQ-MFI of FoxP3 expression in pan- (striped bars) and MOG-specific (open bars) CD4^+^CD25^+^ population in the CNS among young (cherry) and aged mice treated with vehicle (blue) or P300i (black). In panels C and E, each symbol represents an individual animal sample. Data are expressed as mean ± SEM. The *p*-values between three groups were analyzed by one-way ANOVA with a Dunnett’s multiple post-hoc test, and a statistically significant difference was considered to be *p < 0.05,* “N.S.” stands for “not significant”.

These results suggest that the potential underlying mechanism of the mitigation of late-onset EAE severity via the transient inhibition of FoxP3 expression in the accumulated pan-pTreg cells in aged mice is very similar to the mechanism using this method in mitigation of AD pathology ^60^. It is due to easing the cellular trafficking of anti-inflammatory cells ^63^, including not only pan- and MOG-specific Treg cells but likely other anti-inflammatory monocytes, such as monocyte-derived anti-inflammatory M2 macrophages, into the inflamed CNS to enhance the anti-inflammatory capacity.

## DISCUSSION

Typically, human MS disease develops in young (20 – 40 years old) females and presents with at least four clinical sub-types. Two of them, progressive-relapsing MS (PRMS) and RRMS, have relatively mild symptoms, and are often seen in young patients (80% - 90%). However, the other two, PPMS and SPMS, have severe symptoms, and are often seen in aged (>65 years old) patients (~29% for PPMS and ~26% for SPMS), while young patients only account for 10% PPMS and rarely exhibit SPMS ^5^. Late-onset MS in the elderly has been reported ^4, 5, 6, 7^, and the mean age of the MS population is rising ^5^. It was reported that 14% of total MS patients were 65 years and older in 2010 ^64^. The severe symptoms in the aged patients are more or less attributed to Treg cell function, since Treg cells are regarded as protective cells, which combat IFN-g-producing and IL-17-producing CD4^+^ pathogenic cells ^35, 40, 41, 42, 43, 44, 45^, and pathogenic CD8^+^ T cells involved in the MS lesion ^65, 66^.

Ample evidence shows that Treg cells play an ameliorative role in MS/EAE disease onset and severity ^33, 34, 35^. Transferring Treg cells into MOG-induced EAE mice attenuated disease ^67^ and EAE severity correlated inversely with the frequency of MOG-specific Treg cells ^68^. However, accumulation of Treg cells ^25^ with enhanced suppressive function ^26^ in aged individuals is accompanied with severe symptoms in aged mice with late-onset EAE disease (Fig. 1). This is not consistent with the notion that Treg cells play a role in suppressing uncontrolled immune reactions. This inconsistency encouraged us to investigate the underlying mechanism.

CD4^+^FoxP3^+^ Treg cells primarily act to suppress Teff cell-mediated aberrant antigen-specific and non-specific immune responses. Accumulation of Treg cells in the elderly is disadvantageous for fighting infection, cancer, neurodegenerative and autoimmune diseases. In protection and recovery from CNS inflammatory disorders, such as AD, one of the age-related neuroinflammatory diseases, excessive Treg cells were shown to play a detrimental role ^60^. This is probably due to Treg cell distribution either inside the CNS or in the periphery ^1, 2^, in addition to the existence of distinct Treg subsets ^69, 70^. Herein, using an aged EAE mouse model that resembles aged human MS disease, we found that the aged EAE mice had a different distribution of pan- and MOG-specific Treg cells compared to their young counterparts. Specifically, aged mice exhibited a high proportion of pan-Treg and a low proportion of MOG-specific Treg cells in their periphery, but low proportions of both pan- and MOG-specific Treg cells in the inflamed CNS (Figs. 2 and 3). Accumulation of Treg cells outside the CNS and residing at the CNS-periphery boundaries, the BBB and CP, could impede immune cell, including Treg cell, trafficking into the inflamed CNS for the recovery ^60, 63^.

Treg cells are unstable and plastic, and this is especially observable during inflammatory autoimmune stimulation ^16, 51^. The unstable characteristics are exhibited by loss of FoxP3 expression in a proportion of mature Treg cells ^71^, and the plastic characteristics are exhibited by the production of pro-inflammatory cytokines IFN-g or IL-17, along with FoxP3 expression. These Treg cells acquire an effector-like phenotype, and they are able to induce, rather than suppress, autoimmunity. These cells are commonly seen in the autoimmune-prone NOD background mice and diabetic patients ^52, 72^, as well as in MS patients (mouse EAE setting) ^73, 74^ We found that in the aged inflamed CNS Treg cells exhibited higher plasticity than in the young counterparts, observed as co-expression of INF-g and/or IL-17 with FoxP3 (Fig. 4), which potentially results in reduced suppressive function and increased pathology ^16, 53^. This could contribute to the higher clonal expansion of Teff cells in this inflamed CNS (Fig. 5). Furthermore, this also results in disruption of a normal Treg/Teff balance. In addition to self-antigen-driven inflammatory autoimmune stimulation ^16, 51^, the age-related, chronic, systemic inflammation (inflammaging) may play a synergistical role to enhance Treg plasticity in aged EAE mice compared to young counterparts. However, this area needs further investigation.

It is a difficult and complex task to overcome the accumulation of Treg cells in the elderly as a therapeutic strategy for neuroinflammatory disease. Transient inhibition of FoxP3 expression in the accumulated peripheral CD4^+^FoxP3^+^ Treg cells in aged individuals is one option, which was used in AD and displayed improvement of amyloid-beta plaque clearance, amelioration of neuroinflammation, and recovery of cognitive decline demonstrated in an AD mouse model ^60^. We transiently inhibited accumulated peripheral Treg cells in aged mice, which partially ameliorated the EAE disease and corrected Treg distribution (Figs. 6 and 7). The mechanism is probably the accumulated CP-resident Treg cells suppressing INF-g producing cells, such as Th1 CD4^+^ T cells outside the CNS, which directly block the gateway ^60^ for homeostatic leukocyte trafficking into the CNS ^75, 76^. IFN-g is required for activation of the brain’s CP for CNS immune surveillance and repair ^63, 76^. For example, immunization with a myelin-derived antigen was reported to activate the brain’s CP via inducing the CP to express IFN-g and attract Th1 cells, thereby enhancing recruitment of immunoregulatory cells to the CNS to achieve attenuation of neuroinflammatory progression in a mouse model ^77^. Interestingly, a complete depletion of pTreg cells, even though in a transient manner, did not seemingly so efficiently alleviate EAE symptom in the aged mice (Fig. S6C) as partial inhibition of pTreg cells (Fig. 6C), but this undefined result remains to be further investigated.

Although CD4^+^ T cells, both CD4^+^ Teff and CD4^+^FoxP3^+^ Treg cells, are the traditional primary actors in MS/EAE disease pathogenesis and immunoregulation, emerging evidence shows that CD8^+^ T cells also have roles in both immunopathology and immunoregulation, either to exacerbate or mitigate brain inflammation during CNS autoimmunity ^78, 79^. Regarding the MS/EAE immunopathogenesis aspect, myelin-specific CD8^+^ T cells exacerbate brain, though not spinal cord, inflammation via a Fas ligand-dependent mechanism to promote lesion formation in the brain ^80^. In addition, IL-17A secreting CD8^+^ T cells, termed Tc17 cells, in the CNS support Th17 cell-mediated autoimmune encephalomyelitis ^56, 57^. In aged neurogenic niches that comprise neural stem cells, CD8^+^ T cells are increased to inhibit the proliferation of neural stem cells ^55^. This is a disadvantage for the recovery of demyelinating plaques in aged MS/EAE disease. Our results show that IL-17A^+^ CD8^+^ cells were increased in the aged EAE CNS (Fig. 4D), which was consistent with severe symptoms and pathology in aged EAE mice. However, regarding the MS/EAE immunoregulation aspect, although the concept of CD8^+^Treg cells is not unanimously accepted, the function of CD8^+^Treg cells in MS/EAE has received attention ^81, 82, 83^. The main focus of studies in CD8^+^Treg cells in MS/EAE is in young patients or young animals, and there are many unanswered questions regarding how these cells play a role in aged CNS autoimmune inflammation in the elderly. Therefore, the function of these cells in the elderly need to be investigated.

In summary, the results herein provide insights into how accumulated aged polyclonal CD4^+^FoxP3^+^ Treg cells in an inflammatory condition do not ameliorate but are detrimental for CNS repair processes in neuronal inflammation of aged MS demonstrated in the animal model EAE.

## METHODS

### Mice and animal care

C57BL/6 wild-type (WT) mice were used. Aged (18 - 20 months old) mice were ordered from the National Institute on Aging, aged rodent colonies. Control young WT mice were 2 - 3 months old. FoxP3-DTR/GFP mice (Stock No: 016958) were obtained from the Jackson laboratory. All mice were maintained under specific pathogen-free conditions in the animal facilities at the University of North Texas Health Science Center. All animal experiments were performed in compliance with protocols approved by the Institutional Animal Care and Use Committee of the University of North Texas Health Science Center (IACUC-2018-0014, 2021-0020), in accordance with guidelines of the National Institutes of Health.

### Late-onset EAE mouse model and disease score determination

EAE was induced as shown in Fig. 1A. Briefly, MOG_35-55_ peptide (WatsonBio Sciences) was emulsified in Complete Freund’s Adjuvant (Sigma-Aldrich F5881) and given as one-time subcutaneous (s.c.) injection into the upper and lower backs of mice (80μg MOG_35-55_ peptide/10g body weight). Pertussis toxin (PT) (List Biologicals, Cat#179A) was intraperitoneally (i.p.) injected on days 0 and 1 (100ng PT/10g body weight). Mice were monitored daily for EAE symptoms and body weight changes. EAE scores were assigned in supplemental Table S1.

### Transient inhibition of FoxP3 expression in pan-pTreg cells and transient depletion of pTreg cells

As shown in Fig. 6A, 12 days after EAE induction in young and aged WT mice, P300i (C646; Tocris Bioscience, Cat# 4200) was i.p. injected at 8.9 mg/kg body weight per day for 5 consecutive days (5x) in one of the aged mouse groups similar to published protocol ^60^. Vehicle-injected young and aged mice were controls. The dynamic changes of Treg cell frequencies and Foxp3 expression were monitored at three-time points as denoted by red arrows in Figure 6A.

12 days after EAE induction in young and aged DTR mice, DT (Sigma-Aldrich, Cat#D0564) was i.p. injected at 50μg/kg body weight to one of the aged-DTR mouse groups. Vehicle-injected young and aged mice were controls.

### Cell isolation from the CNS and the CP

For mononuclear cell isolation from the brains and the spinal cords, euthanized mice were transcardially perfused with 20ml of PBS. CNS tissues were minced and digested with 2mg/ml collagenase D (Roche, Cat# 11088858001) and 28U/ml DNase I (Invitrogen, Cat# 18068015) in RPMI-1640 at 37 °C for 45 min, followed by percoll (Sigma-Aldrich, Cat# P1644) gradient centrifugation per our previous method ^84^. For cell isolation from the brain CP, CP tissues (5 mice per group) were collected from the lateral, third and fourth ventricles of the brain, then digested at 37°C for 40 min in 2mg/ml collagenase-D/PBS with pipetting.

### Tetramer-based flow cytometric assay of pan- and MOG-specific Treg and Teff cells and IFN-γproducing cells at the CP

Single-cell suspensions from the LN, spleen, and the CNS of mice were stained with extracellular fluorochrome-conjugated CD markers (Biolegend), along with APC-conjugated MOG38-49 I-A^b^ tetramer (NIH tetramer core, peptide sequence: GWYRSPFSRVVH) and Brilliant Violet 421-conjugated human CLIP_87-101_ I-A^b^ tetramer as a control (peptide sequence: PVSKMRMATPLLMQA). Then, cells were fixed and permeabilized using the fixation and permeabilization kit (eBioscience, Cat# 00-5523-00) for intracellular staining of PE-conjugated anti-FoxP3 (eBioscience Cat# 12-5773-82) per the company’s instruction. Data were analyzed using FlowJo™ v10 software and particularly, MFIs were defined as the medians of fluorescence intensities of the conjugated fluorochromes of the antibodies.

Cells isolated from the CP were incubated with PMA (5ng/ml), ionomycin (500ng/ml), and Protein Transport Inhibitor (0.7μL/mL, BD Biosciences. Cat# 51-2092KZ) for 5hrs, followed by extracellular staining with PerCP/Cy5.5 conjugated anti-CD45 (Biolegend, Cat# 103132) and intracellular staining with APC-conjugated anti-IFN-g (Biolegend, Cat# 505810).

### Single-cell RNA-seq of CNS T cells for analysis of transcriptome profile and TCR copy number-based T cell clonal expansion

Mononuclear cells were isolated from the CNS of three young and three aged EAE mice. Cells were stained with anti-CD3-APC (Biolegend, Cat#100236) and anti-CD11b-Brilliant Violet 711 (Biolegend, Cat#101242), then sorted on Sony SH800 Cell Sorter to collect CD3^+^CD 11b^-neg^ T cells, followed by gene expression (GEX) and TCR V(D)J library preparation and sequencing with the 10x Genomics Chromium Single Cell 5’ GEM, Library & Gel Bead Kit v2 (10x Genomics, Cat#1000287). The cDNA was amplified using the same kit. Products were purified using Ampure XP beads and quality was controlled using Agilent Tapestation and Qubit 4 fluorometer. TCR target enrichment was done by Chromium Single Cell Mouse TCR Amplification kit (10x Genomics, Cat #1000254). TCR V(D)J and GEX libraries were constructed by the Library Construction Kit (10x Genomics, Cat #1000190) with Dual Index Kit TT Set A. Sequencing was on an Illumina NovaSeq 6000 according to 10x Genomics sequencing protocol recommendations.

Fastq files (Cell Ranger, version 6.0.2, 10xGenomics provided mm10 reference genome), Cloupe file, and Vloupe files were generated for the downstream analysis. The t-SNE plots of T cells were visualized by 10x Genomics Loupe Brower 5.0 and T cells were classified into three groups, including CD8 T cells, CD4 Teff cells and CD4 Treg cells. Numbers of unique molecular identifiers (UMI) of *Ifng* and *Il17a* were used to determine the gene expression levels in every single cell in the Treg and CD8^+^ T cell populations. 10x Genomics Loupe VDJ Brower 4.0 was used to output the clonotypes of CD4^+^ Teff and Treg cells by aggregating the Cloupe file and Vloupe file of each sample. Each clone was determined by the CDR3 regions of paired TCRa and TCRβ chains. Based on TCR sequence similarity, clones with clonal size greater than 2 were defined as expanded clones, among which the most frequent 10 clones in each sample were defined as the top 10 expanded clones. All unique clones were defined as unexpanded clones.

### Luxol Fast Blue staining for demyelination assay of spinal cords

Paraffin sections of the spinal cord (Fig. 6D) were stained with 0.1% Luxol Fast Blue (LFB) solution (Sigma-Aldrich, Cat#S3382) per a previous publication ^85^, with a modification adding eosin counterstaining (details in Supplemental Method-B).

### Statistics

Statistical tests used to analyze each set of experiments are indicated each figure legend, including the unpaired two-tailed Student’s *t*-test for two groups, and one-way ANOVA for multiple groups. Data from EAE scoring were analyzed by Mann-Whitney *U* test to compare between two groups, and Kruskal-Wallis test was used to compare multiple groups followed by Dunnett’s multiple comparisons post hoc test for pairwise comparisons of groups. Body weight changes along with time between two groups were analyzed with two-way repeated-measures ANOVA with Geisser-Greenhouse correction. Results were considered statistically significant at values of \**p* < 0.05; \*\**p* < 0.01; \*\*\**p* < 0.001; **** *p* < 0.0001.

### Data Availability

All scRNA-seq raw data are available on the Gene Expression omnibus (GEO) (GSE182747) [https://www.ncbi.nlm.nih.gov/geo/query/acc.cgi?acc=GSE182747]. All individual numerical values, detailed statistic information and other regarding source data in each figure are included in source data Excel file.

## Acknowledgements

We thank Drs. Rance Berg and Sterling Ortega (UNTHSC) for critiques of experiments and discussion, Dr. Nicky Hales (previously UNTHSC, current affiliation: 10x Genomics) for performing single cell immune profiling, and the National Institutes of Health Tetramer Core Facility at Emory University for providing the tetramer reagent.

## Author contribution

W.W. performed most of the experiments and hands-on animal work, as well as wrote the manuscript; R.T. assisted with cell sorting experiments and edited the manuscript; J.O. performed some experiments; D-M.S. conceived and designed project, did some hands-on animal work, analyzed data, and wrote the manuscript.

## Abbreviations

AD: Alzheimer’s disease
Ag: antigen
BBB: blood-brain barrier
CP: choroid plexus
CNS: central nervous system
DPI: days post-immunization
DT: diphtheria toxin
DTR: diphtheria toxin receptor
EAE: experimental autoimmune encephalomyelitis
LN: lymph nodes
MFI: mean florescence intensity
MOG: myelin oligodendrocyte glycoprotein
MS: multiple sclerosis
pan-: polyclonal
PPMS: primary progressive MS
PRMS: progressive relapsing MS
RRMS: relapsing-remitting MS
SPMS: secondary progressive MS
PT: Pertussis toxin
pTreg or tTreg: peripheral Treg or thymic Treg
sc-RNA-Seq: single-cell RNA sequencing
Teff: T effector cell
Treg: regulatory T cell

---

**Supplemental Figure S1.**
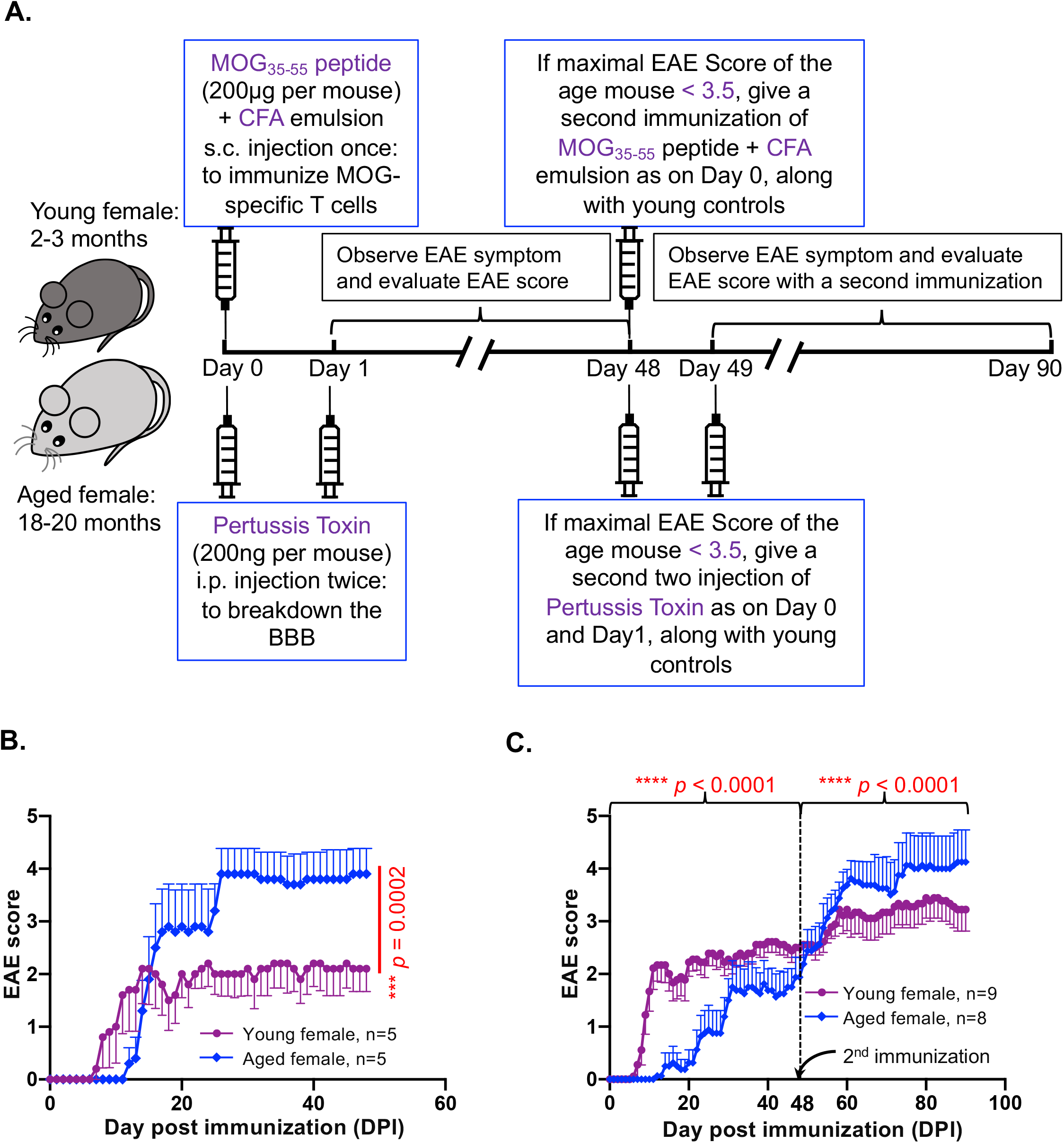
Two types of EAE scores of aged female mice with equal dosage of MOG_35-55_ peptide as young mice. **(A)** Workflow for immunization and induction of EAE in C57BL/6 mice with MOG peptide (200μg per mouse) and EAE disease course observation. **(B)** Type-I EAE disease course in aged female mice with young female mice as control. **(C)** Type-II EAE disease course in aged female mice with young female mice as control. The p-values of EAE scores were calculated by Mann-Whitney U test and a statistically significant difference was considered to be *p* < 0.05.

**Supplemental Figure S2.**
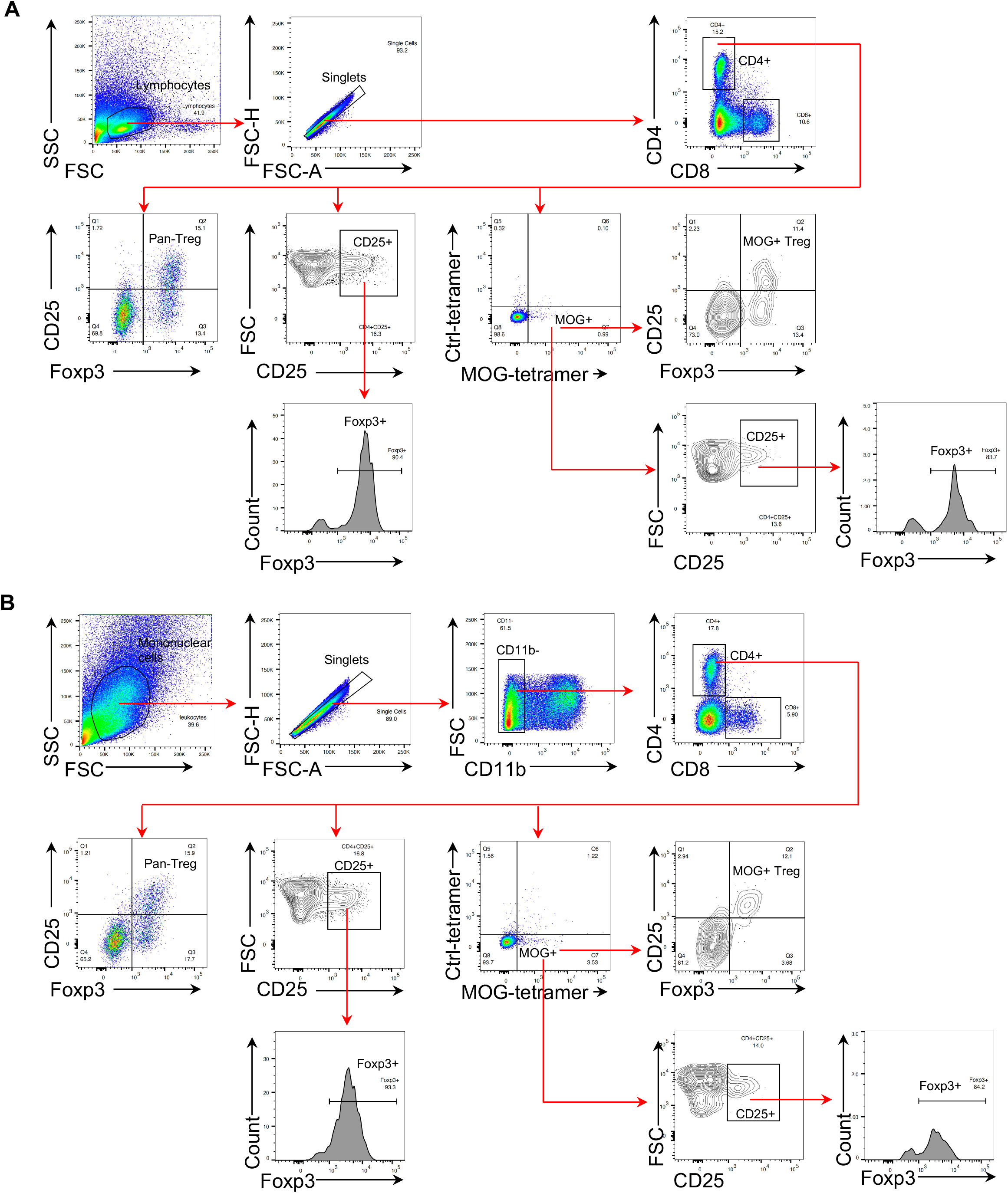
Flow cytometric gating strategies of T cell populations in the periphery and the CNS. **(A)** representative gating strategy for the T cell populations in the periphery (spleen/lymph node). **(B)** representative gating strategy for the T cell populations in the CNS (brain/spinal cord).

**Supplemental Figure S3.**
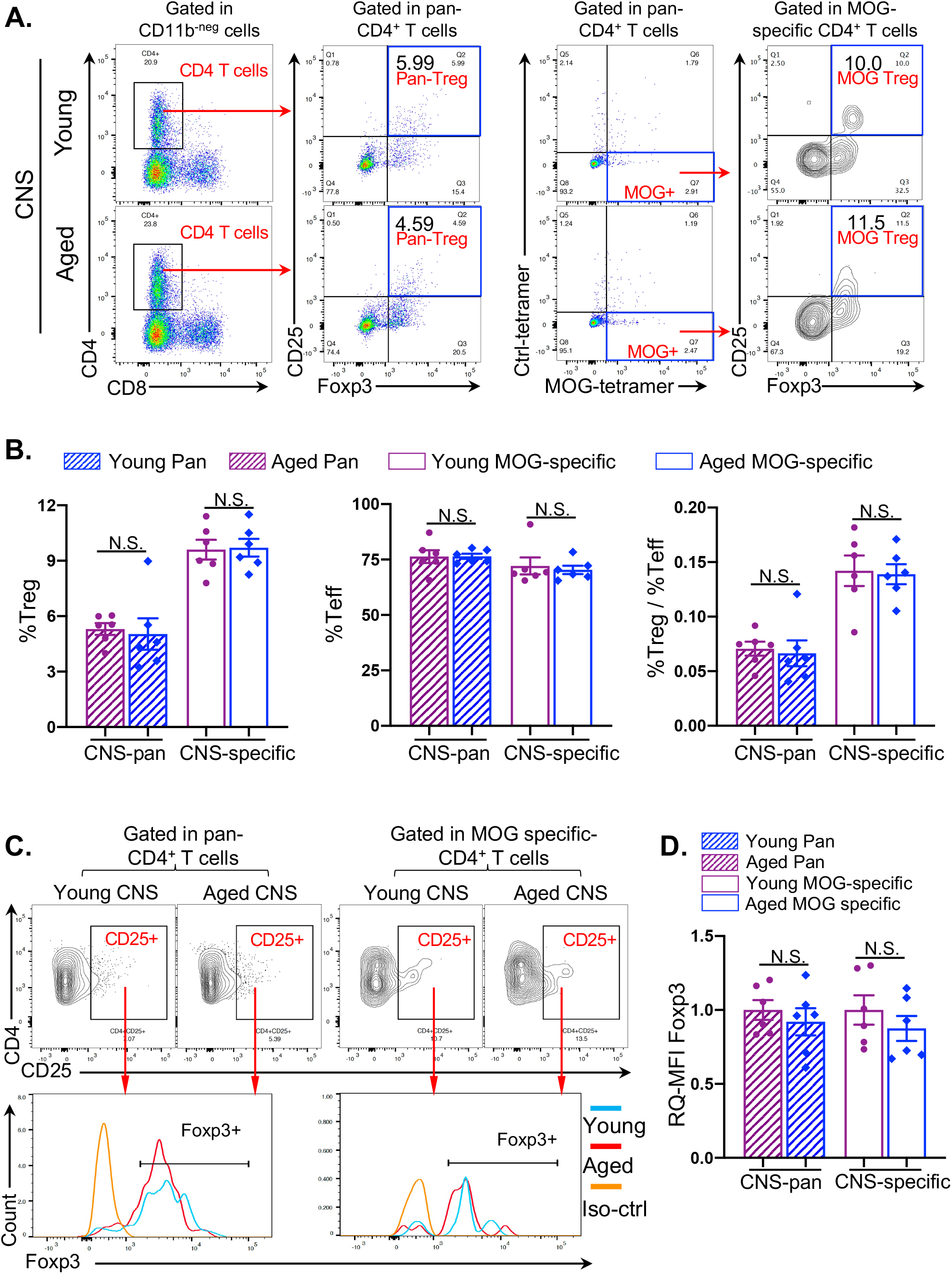
Balanced pan- and MOG-specific Treg distributions in the CNS of late-onset EAE disease (8 days after immunization with MOG) in aged mice. **(A)** Flow cytometry gating strategies show pan-Treg population (left two columns) and MOG-specific Treg population (right two columns) in the CNS. **(B)** Summarized results of the percentages of pan- (striped bars) and MOG-specific (open bars) Treg cells (top panel) and Teff cells (middle panel), and ratios of Treg/Teff cells (bottom panel) in the CNS (brain and spinal cord) between young (cherry) and aged (blue) mice. **(C)** Flow cytometry gating strategies show representative FoxP3+ gates (bottom panels) from CD4^+^CD25^+^ gates of the CNS (mix of brain and spinal cord) of young and aged mice. **(D)** Summarized results of the RQ-MFI of FoxP3 expression in pan- (striped bars) and MOG-specific (open bars) CD4^+^CD25^+^ population in the CNS between young (cherry) and aged (blue) mice. In panels B and D, each symbol represents an individual animal sample. The statistical significance between the two groups was analyzed by unpaired Student’s *t*-test, “N.S.” stands for “not significant”, and error bars indicate mean ± SEM.

**Supplemental Figure. S4.**
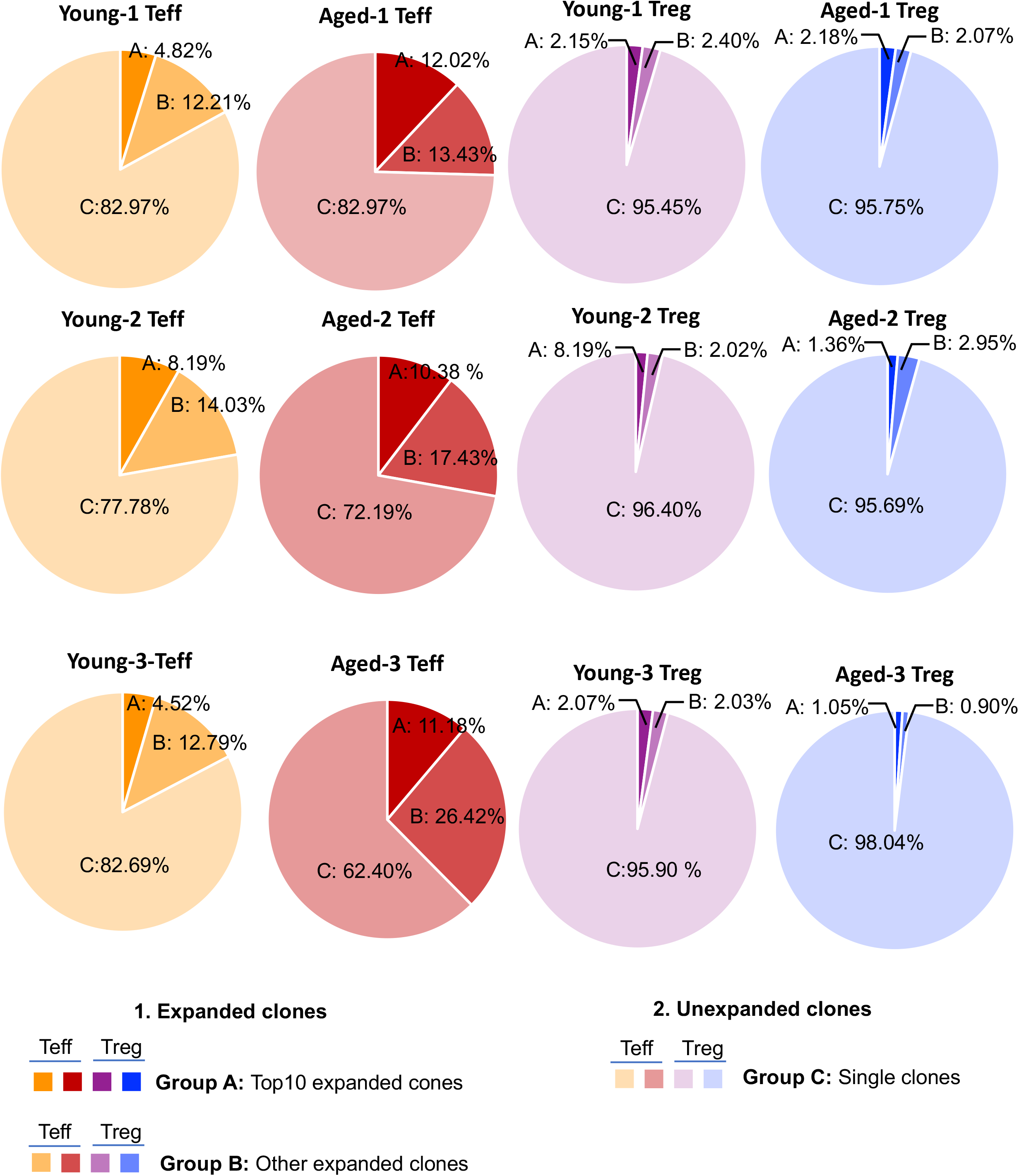
Clonal expansion in CNS-infiltrated CD4+ T populations of individual young and aged mice during EAE. Leftmost column: individual pie charts of young Teff cell clonal expansion; second left column: individual pie charts of aged Teff cell clonal expansion; second right column: individual pie charts of young Treg cell clonal expansion; rightmost column: individual pie charts of aged Treg cell clonal expansion.

**Supplemental Figure S5.**
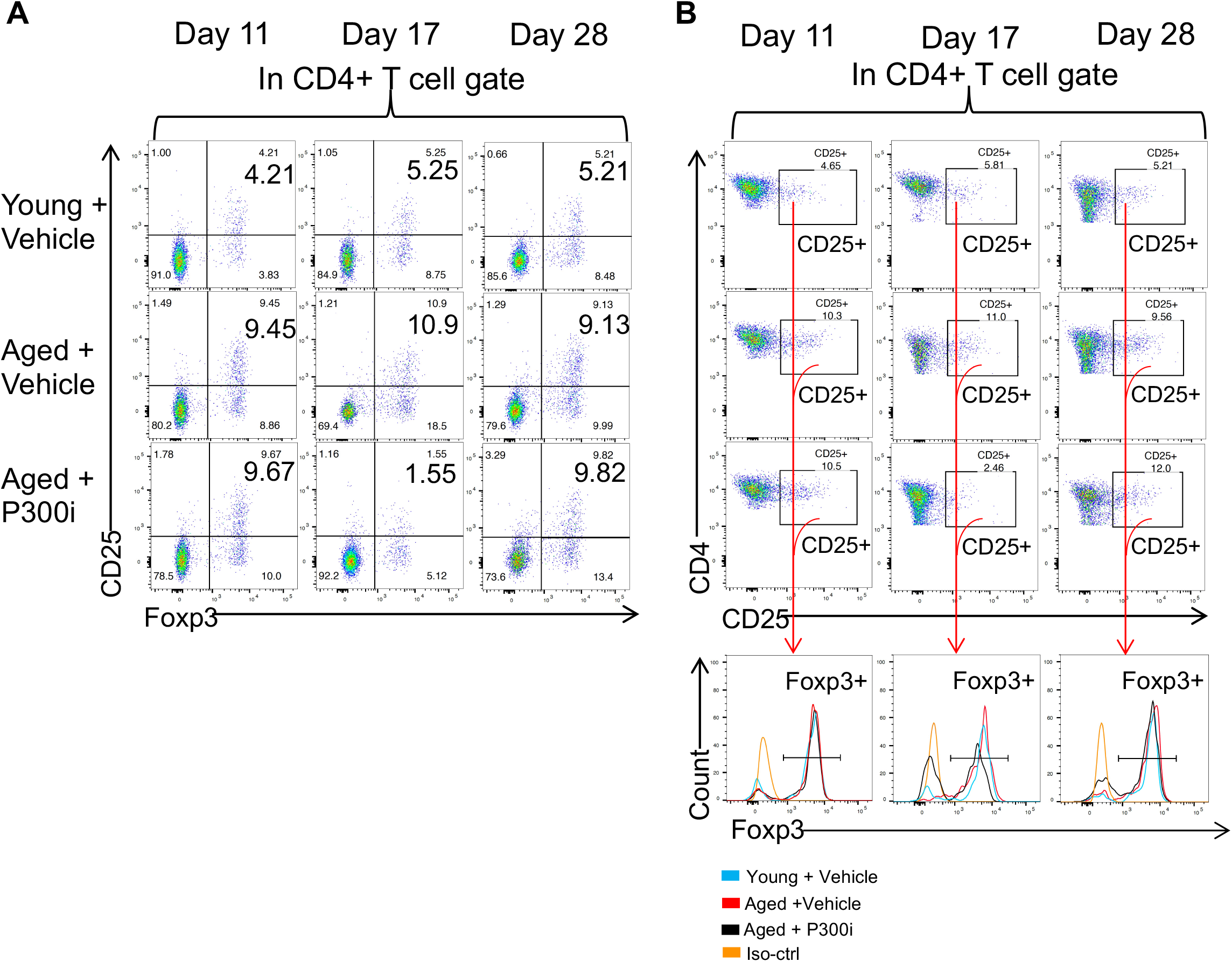
Gating strategies of peripheral blood Treg cell dynamics during partial and transient pTreg cell inhibition. **(A)** Flow cytometry gating strategies show the frequencies of peripheral blood Treg cells in the three groups of mice 11-DPI (one day before p300i or vehicle treatment), 17-DPI (one day after the last p300i or vehicle treatment) and 28-DPI (12 days after the last p300i or vehicle treatment). **(B)** Flow cytometry gating strategies show representative FoxP3+ gates (bottom panels) from CD4^+^CD25^+^ gates of the peripheral blood Treg cells in the three groups of mice 11-DPI, 17-DPI, and 28-DPI.

**Supplemental Figure. S6.**
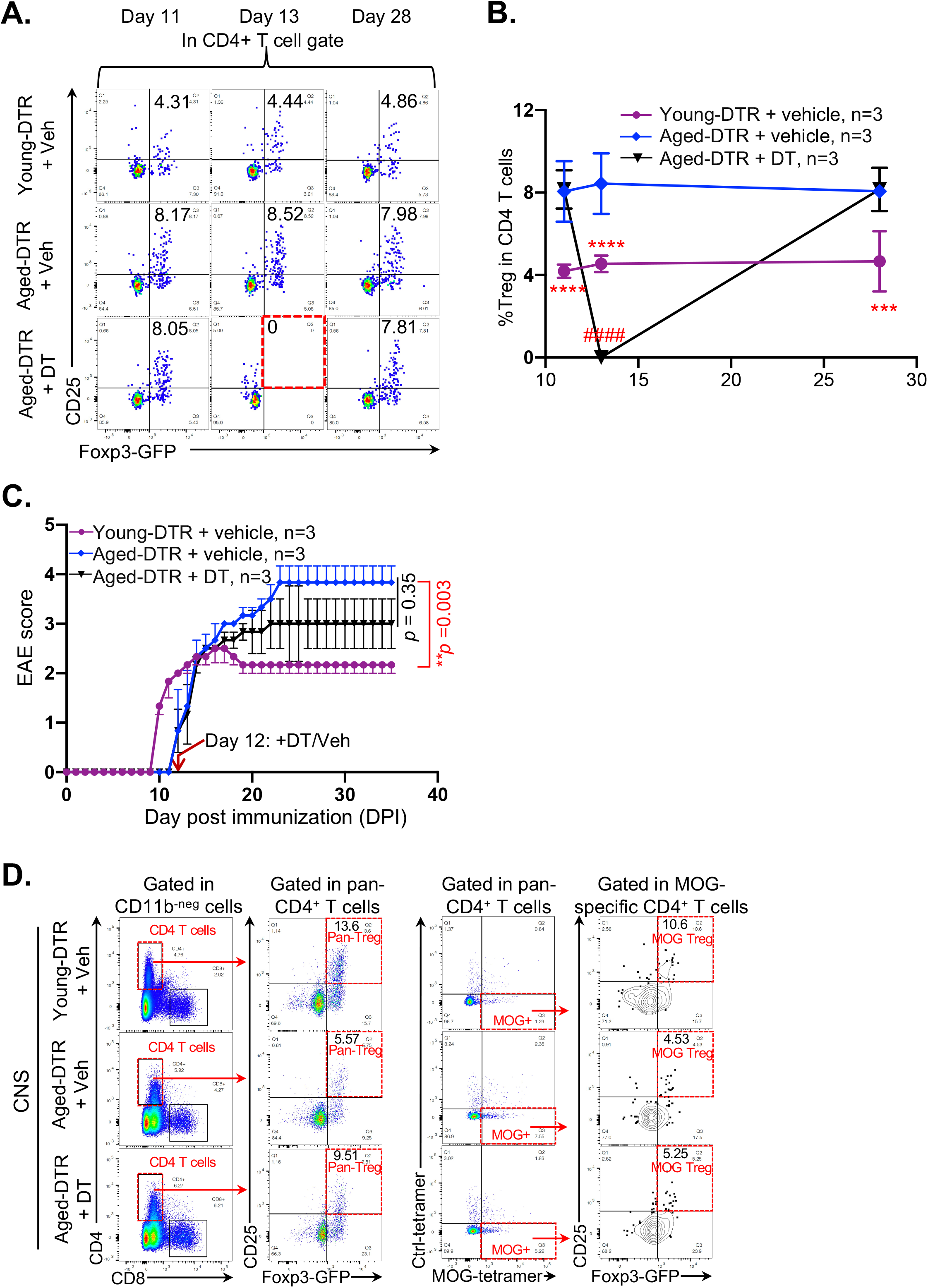
Alleviation of EAE severity after transient depletion of accumulated pTreg cells in late-onset EAE in aged Foxp3-DTR mice. **(A)** Flow cytometry gating strategies show the frequencies of peripheral blood Treg cells in the three groups of mice 11-DPI (one day before DT or vehicle treatment), 13-DPI (one day after DT or vehicle treatment) and 28-DPI (16 days after DT or vehicle treatment). **(B)** Summarized results of dynamic changes of Treg frequencies in the peripheral blood of three groups of mice. All data are expressed as mean ± SEM and are analyzed by One-way ANOVA followed by Dunnett’s multiple post-hoc test. *** *p* < 0.001, young-DTR mice + vehicle v.s. Aged-DTR mice + vehicle; **** *p* < 0.0001, young-DTR mice +vehicle v.s. Aged-DTR mice +vehicle; #### *p* < 0.0001, aged-DTR mice + vehicle v.s. aged-DTR mice + DT. Animal numbers in each group are listed in each panel. **(C)** Alleviation of the symptoms in late-onset EAE of aged-DTR mice. The *p*-values were calculated with Kruskal-Wallis test followed by Dunnett’s multiple comparisons post hoc test for pairwise comparisons of groups, with p < 0.05 as statistically significant. **(D)** Flow cytometry panTreg and MOG-specific Treg cells in the CNS of young-DTR and aged-DTR mice with DT or vehicle treatment.

## Supplemental Methods

### A.

**Table S1:** Mouse EAE scoring

A. Table S1: Mouse EAE scoring
| Score | Clinic observations |
| --- | --- |
| 0 | No obvious changes in motor function compared to non-immunized mice |
| 0.5 | tail tip limpness |
| 1.0 | tail limpness |
| 1.5 | tail limpness and hind leg inhibition |
| 2.0 | partial hind leg paralysis |
| 2.5 | partial hind leg paralysis with dragging of at least one hind leg |
| 3.0 | complete paralysis of hind legs |
| 3.5 | complete hind leg paralysis and unable to right the body when placed on the side |
| 4.0 | complete hind leg and partial front leg paralysis |
| 4.5 | complete hind leg and partial front leg paralysis and mouse is not alert |
| 5.0 | spontaneously rolling in the cage or death |

### B. Luxol fast blue (LFB) staining * of demyelinated spinal cord with/without Eosin counterstaining

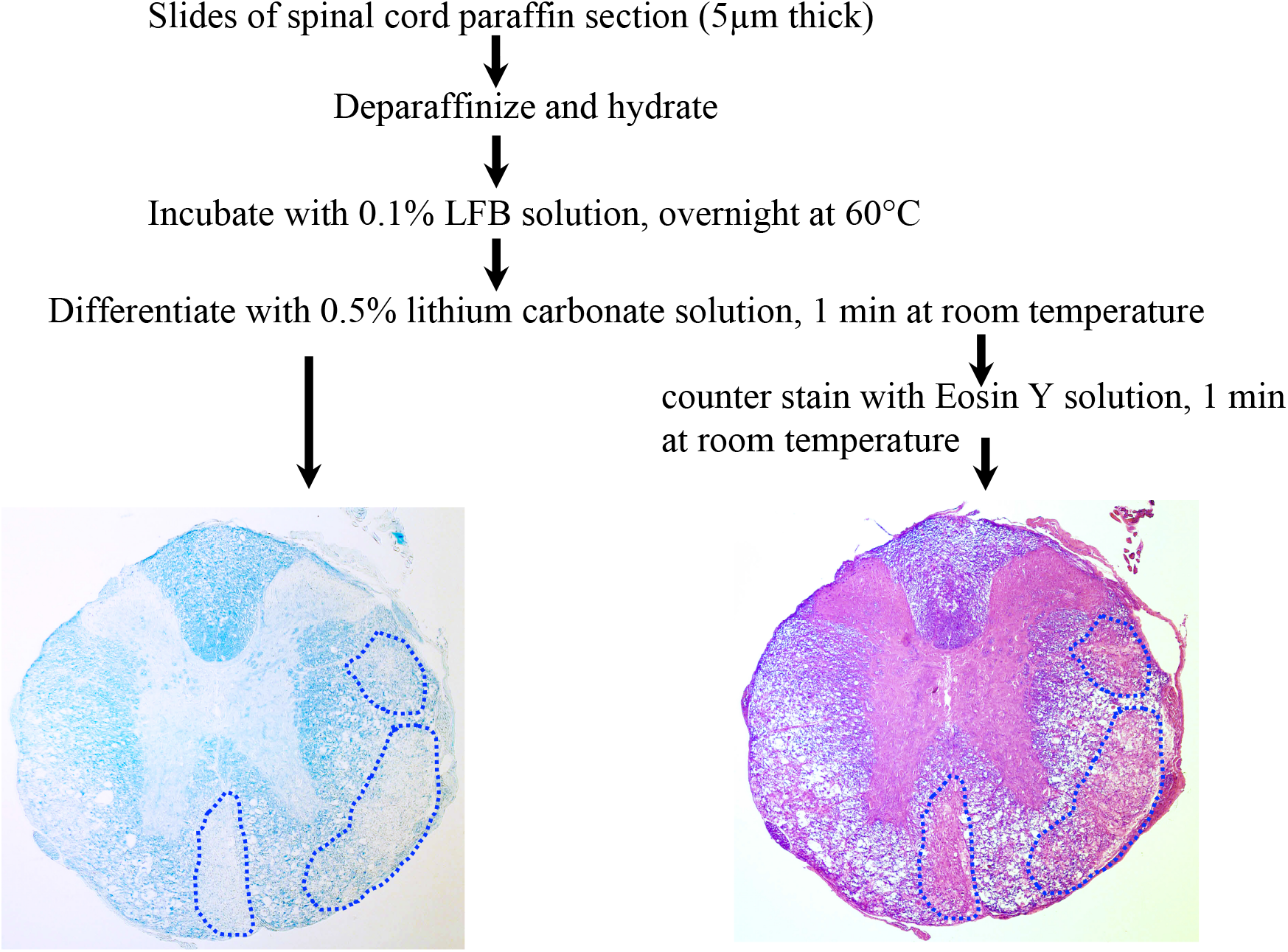

